# A Novel Eukaryotic Ribosome Factor Enables Translation Restart Following Cellular Dormancy

**DOI:** 10.64898/2026.03.21.713407

**Authors:** Maciej Gluc, Higor Rosa, Maria Bozko, Lesley A Turner, Cassidy R Prince, Yelena Peskova, Heather A Feaga, Kathleen L Gould, Simone Mattei, Ahmad Jomaa

**Author notes:** These authors have contributed equally to this work (Maciej Gluc, Higor Rosa).

## Abstract

Dormancy is a survival strategy employed by all domains of life to withstand prolonged nutrient deprivation and environmental stress that is marked by a global shutdown of protein synthesis. However, the molecular mechanisms driving ribosome inactivation and reactivation during and after dormancy in eukaryotes remain poorly understood. Here, we identify SNOR, a novel SBDS-like ribosome-associated factor in Schizosaccharomyces pombe, that is upregulated and associates with ribosomes during induced dormancy triggered by glucose depletion. SNOR contributes to protein synthesis repression by binding the ribosome to probe the peptidyl transferase center (PTC), block tRNA-binding sites, and cap the polypeptide exit tunnel (PET). Importantly, we show that SNOR is essential for the restart of protein synthesis upon glucose reintroduction and exit from dormancy. SNOR is evolutionarily conserved and specifically upregulated in response to glucose stress in fungi. These findings reveal a previously unrecognized ribosome-associated factor that links glucose stress and cellular dormancy to surveillance of protein synthesis and highlight the power of in situ structural biology to uncover stress-responsive regulators of translation.

## Introduction

Nutrient deprivation triggers metabolic stress pathways and initiates adaptive programs to sustain cell survival. Under prolonged stress, cells can enter a reversible, non-proliferative dormant state characterized by low metabolic activity that allows them to survive extended periods of nutrient or water deprivation^1,2^. Dormancy is observed across diverse eukaryotic systems, including fungi, protozoa, plants, animals, and even cancer cells^3–6^. In fungi, dormant cells can withstand oxidative and pH stress due to harsh environmental conditions, and in some cases, can persist in host tissues by evading immune response for many years before reactivating^7^. Importantly, slow-growing or dormant cells formed by pathogenic fungi are also less susceptible to antifungal drugs and contribute to drug treatment failure, persistence, and infection relapse^8^.

During exponential growth, yeast can devote up to 50% of cellular energy to protein synthesis. This process is rapidly downregulated under unfavorable metabolic conditions that induce cellular dormancy^9^. Ribosome shutdown is an integral part of downregulating protein synthesis and is mediated by a specialized group of proteins, known as hibernation factors ^10–13^. These factors engage with critical ribosomal sites including the tRNA binding pockets, mRNA entry channel, PTC and polypeptide exit tunnel. Through this extensive interaction network, hibernation factors not only block translation but also protect ribosomes from nuclease-mediated degradation^14–16^. Upon return to favorable conditions, these factors dissociate or are displaced, enabling translation to resume and restoring protein synthesis^17,18^.

The first ribosome hibernation factors were discovered over 30 years ago in bacteria, including the ribosome modulation factor (RMF) and hibernation promoting factor (HPF)^14,19,20^. Since then, functionally similar yet structurally and evolutionarily distinct proteins have also been identified in eukaryotes, particularly in the context of development, stress responses, and cellular quiescence^15,16,21–23^. Despite their common function, these factors exhibit remarkable mechanistic and structural diversity. Within a single organism, multiple hibernation factors can respond to specific environmental stressors, suggesting that varied pathways for ribosome hibernation are important for prolonged survival ^12,13^. Altogether, these discoveries point to a broader and more functionally diverse landscape of ribosome regulation than what is currently known.

Advances in *in-situ* cryo-ET have allowed us to determine how various factors act in concert to orchestrate ribosome hibernation. Using this approach, we identified SNOR as a protein that monitors the PTC to reduce translation in fungi undergoing glucose limitation. Upon access to fresh nutrients, SNOR is important for translational restart, likely acting together with the universally conserved translation factor eIF5A. Therefore, our results provide essential insight into a previously unknown mechanism of translational restart after dormancy exit in living cells.

## Results

### In situ cryo-ET reveals an SBDS-like protein bound to the PTC of hibernating ribosomes

Our previous cellular cryo-ET study of glucose depleted *Schizosacharomyces pombe* revealed the presence of outer mitochondrial membrane (OMM)-associated ribosomes in dormant cells (Fig. 1a). The *in situ* structural analysis showed that OMM-tethered ribosomes were in a hibernating state bound to elongation factor eEF2 and devoid of tRNA^24^. EM-densities were also visible at the ribosome subunit interface within the tRNA binding sites, suggesting the potential presence of additional hibernation factors. However, the limited resolution of the map (∼11 Å) hindered our ability to identify these factors.

**Figure 1.**
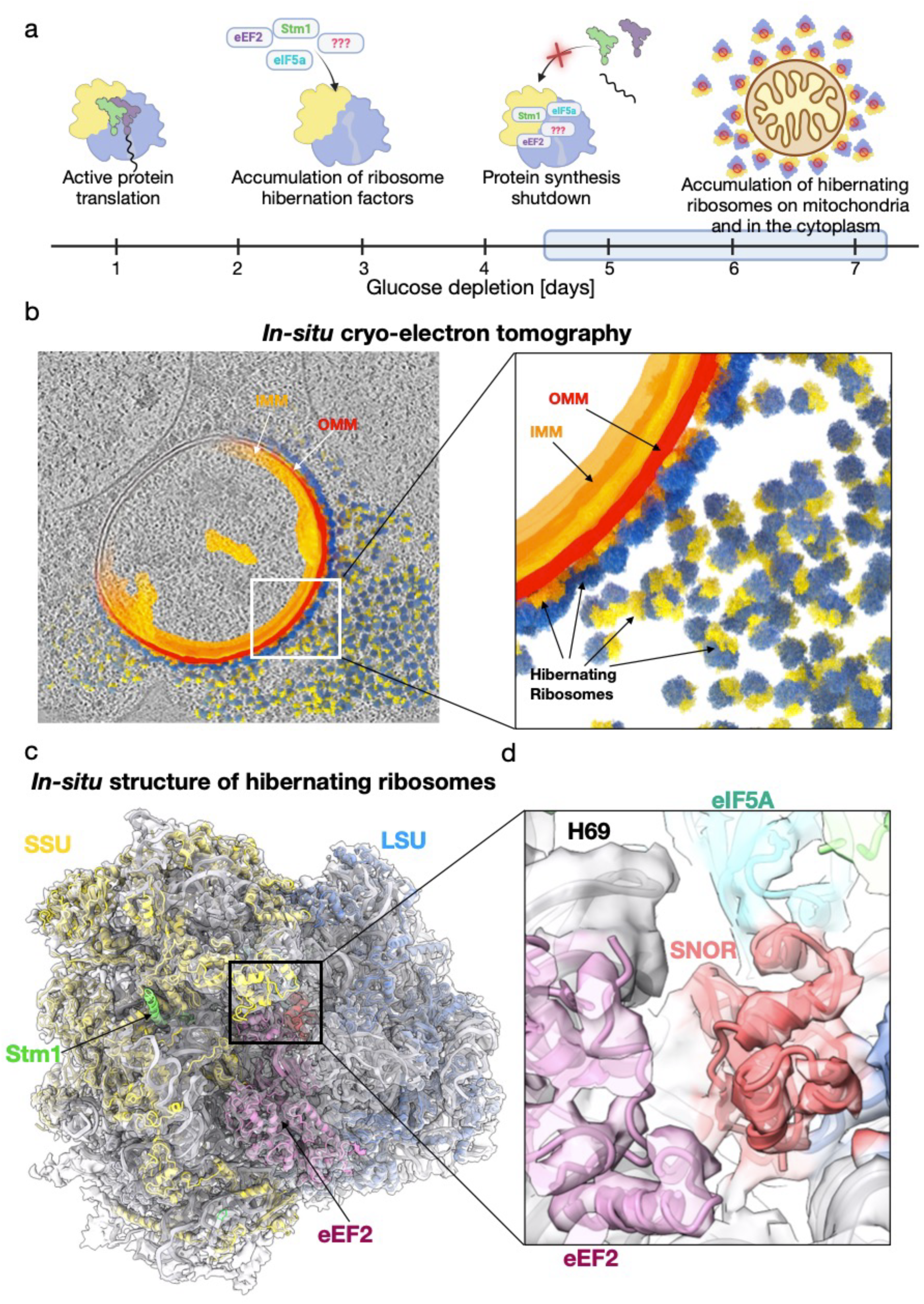
Identification of SNOR, an SBDS-like factor bound to hibernating ribosomes by *in situ* cryo-ET. (a) Schematic representation of prolonged glucose depletion effects on *S. pombe* ribosomes. The small subunit (SSU) is shown in yellow, the large subunit (LSU) in blue, and the polypeptide exit tunnel (PET) in gray. (b) Computational slice through a tomogram of a mitochondrion decorated with ribosomes from a *S. pombe* cell grown for 7 days in EMM containing 0.5% (w/v) glucose. Partial segmentation highlights the mitochondrion and surrounding ribosomes. Inset shows a close-up of ribosomes at the mitochondrial surface; the inner mitochondrial membrane (IMM) is shown in orange, the outer mitochondrial membrane (OMM) in red, and ribosomal SSU and LSU in yellow and blue, respectively. (c) Cryo-ET reconstruction of the hibernating ribosome at 5.5 Å resolution, shown as a semi-transparent surface with the fitted atomic model rendered as a cartoon. Ribosomal proteins of the SSU and LSU are colored yellow and blue, respectively. Stm1 is shown in green, eEF2 in purple, eIF5A in cyan, and SNOR in red. (d) Inset shows a close-up of SNOR interface at the peptidyl transferase center (PTC).

To improve the quality of the *in-situ* map, we further optimized our sample preparation, data collection, and processing workflows. First, we prepared 32 cellular lamellae by focused ion beam milling at cryogenic temperature (cryo-FIB) with nominal thickness of 150 nm, followed by manual polishing. A total of 175 tilt series were collected from 17 lamellae and 78 reconstructed tomograms with sample thickness lower than 200 nm were selected for image processing. Using the previously determined map of cytosolic hibernating ribosomes^24^ as template, we performed template matching to identify and extract a total of 43,955 ribosomal particles for structural determination, followed by rounds of refinement and classification. To investigate whether the structure of hibernating ribosomes tethered to mitochondria differs from those freely localized in the cytosol (Fig. 1b; Extended Data Fig. 1) we aligned and averaged 12,174 OMM-associated ribosomes and 14,006 unbound ribosomes, yielding reconstructions at 6.0 Å resolution and 6.6 Å resolution, respectively. Structural alignment of the two cryo-ET maps revealed no discernible differences and confirmed that both ribosome populations were characterized by similar unassigned EM-densities, possibly corresponding to hibernation factors. Therefore, we combined the two datasets to increase the total number of particles contributing to the ribosome reconstruction, resulting in an improved map with an average resolution of 5.5 Å and local resolution of the large subunit (LSU) extending to 4.7 Å (Fig. 1c; Extended Data Fig. 1 and Extended Data Table 1).

The *in situ* structure of the hibernating ribosome revealed a complex network of factors bound to the ribosome that act in concert to maintain its inactive, hibernating state. In addition to the previously identified elongation factor eEF2^24^, we observed several well-established ribosome-associated factors, including the hibernation factor Oga1, the ortholog of *S. cerevisiae* Stm1. Notably, we also detected density corresponding to the initiation factor eIF5A (Extended Data Fig. 2), which was recently reported in hibernating ribosomes isolated from zebrafish eggs^21^.

We identified an additional electron density corresponding to a small globular domain located between eEF2 and eIF5A, near helix 69 (H69) covering the PTC of the ribosome. Based on its location and the observed fold of the putative hibernation factor, we screened several candidate factors known to interact with the ribosome at this site. Surprisingly, the best fit into the EM-density was obtained with a ribosome biogenesis factor known as Sdo1, the *S. pombe* homolog of human SBDS. During late stages of assembly, Sdo1 binding is required for the release of anti-association factor eIF6 from the 60S ribosomal subunit^25^. Initial docking of the atomic model of Sdo1 into our map revealed that only its N-terminal SBDS domain (domain I) fits well with the observed density (Extended Data Fig. 3a). No corresponding densities were observed for domains II and III. Furthermore, domain III would clash sterically with the nearby eEF2 density observed in the current structure (Extended Data Fig. 3b).

Given the strong correspondence between the observed density and the N-terminal domain of Sdo1, we hypothesized that this region might represent a distinct factor homologous to the N-terminal SBDS domain of Sdo1. To test this, we first performed a BLAST search of *S. pombe* protein databases using the amino acid sequence of the N-terminal domain Sdo1. However, this search only identified hits of Sdo1 from different organisms, and limiting the search to *S. pombe* did not identify any new proteins. We next turned to Foldseek ^26^, a web-based tool that identifies structural homologs by aligning query models to protein structure databases. Using the atomic model of the truncated N-terminal domain of Sdo1 as a query, we identified a protein of unknown function, Rtc3 (YHR087W). An ortholog of this protein exists in *Saccharomyces cerevisiae,* which has proposed roles in RNA metabolism and in translation ^27–30^. Due to its proposed new function in cell dormancy that we demonstrate, this factor is referred to here as **<u>S</u>**BDS-domain containing hibernatio**<u>N</u>**fact**<u>OR</u>**, or SNOR (Fig. 1d).

### SNOR expression is induced by glucose depletion and specifically binds the ribosome

We first investigated SNOR expression levels during glucose depletion and other stress-related conditions associated with nutrient deprivation. We performed reverse transcription quantitative PCR (RT-qPCR) to monitor changes in mRNA levels during progressive glucose depletion. Samples were collected on Days 1, 2, and 3 of glucose depletion course in Edinburgh Minimal Medium (EMM) containing 0.5% glucose, based on prior observations indicating that global protein synthesis is markedly repressed by Day 4^24^. Transcripts for the genes *eft201, tif51, oga1* and *rtc3* encoding eEF2, eIF5A, Stm1, and SNOR, respectively, were elevated after 3 days of glucose deprivation, with *rtc3* transcript upregulation being the most prominent among the analyzed hibernation factors (Fig. 2a, Extended Data Table 2). To assess whether SNOR protein levels were similarly upregulated, we used CRISPR-Cas9 to introduce a 2×FLAG epitope at the endogenous SNOR locus, generating an N-terminally tagged SNOR protein. Immunoblotting with anti-FLAG over a 7-day glucose depletion course in EMM containing 0.5% glucose showed a significant increase in SNOR protein levels at Days 4 and 7 of glucose deprivation (Fig. 2b).

**Figure 2.**
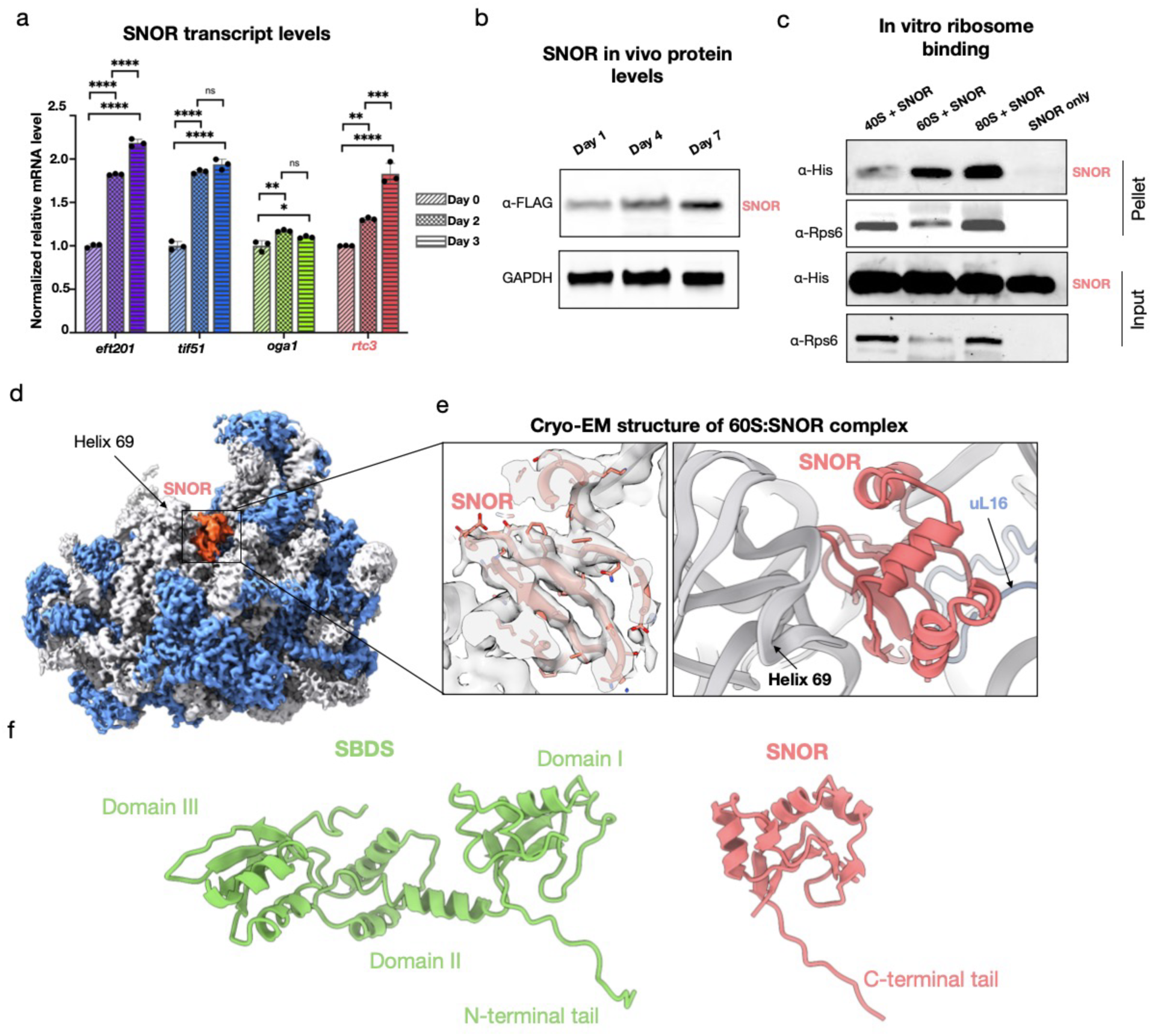
SNOR expression increases during glucose depletion and binds ribosomes. (a) Normalized relative mRNA levels of *eft201* (eEF2, purple), *tif51* (eIF5A, blue), *oga1* (Stm1, green), and *rtc3* (SNOR, red) at days 1 (diagonal stripes), 2 (checkered), and 3 (horizontal stripes) of glucose depletion (n = 3 biologically independent samples). Data are shown as mean ± s.d.. Statistical analysis was performed using one-way ANOVA. *P < 0.05, **P < 0.01, ***P < 0.001, ****P < 0.0001. (b) Immunoblot analysis of SNOR levels in *S. pombe* cell lysates collected after 1, 4, and 7 days of glucose depletion. SNOR was detected using anti-FLAG antibody; GAPDH served as a loading control. (c) Immunoblot analysis of SNOR binding to 40S, 60S, and 80S ribosomal subunits using co-sedimentation assays. SNOR was detected via His-tag; Rps6 was used as a loading control. (d) Cryo-EM reconstruction of the *in vitro* reconstituted complex between the ribosomal large subunit and SNOR. Ribosomal proteins are shown in blue, rRNA in gray, and SNOR density in coral. (e) Close-up view of the atomic model showing SNOR bound to the peptidyl transferase center (PTC) of the ribosome. (f) Structural comparison of an AlphaFold model of SBDS/Sdo1 (left, green) and SNOR (right, coral) atomic models.

To determine whether SNOR directly associates with the ribosome *in vitro*, we recombinantly expressed and purified SNOR from *E. coli* and conducted binding assays using purified *S. pombe* ribosomal subunits. SNOR was incubated with either 80S ribosomes, 60S large subunits, or 40S small subunits, followed by ribosome co-sedimentation using sucrose cushion ultra-centrifugation. The binding assays showed that SNOR specifically interacts with the 60S large subunit and intact 80S ribosomes (Fig. 2c). To further elucidate how SNOR engages the ribosome and describe the molecular interactions responsible for stabilizing its binding to rRNA, we determined the structure of the *in vitro* reconstituted complex between the 60S ribosomal subunit and SNOR (60S:SNOR complex) using single-particle cryo-electron microscopy (SPA cryo-EM). Given that SNOR binds both 60S and 80S ribosomes, we focused on the 60S subunit to minimize sample heterogeneity arising from intersubunit rotational dynamics, thereby facilitating cryo-EM image processing. The final structure of the 60S:SNOR complex was resolved at a global resolution of 2.9 Å, with local resolution for SNOR ranging from 3.5 to 5 Å (Extended Data Fig. 4). The SPA cryo-EM map of the *in vitro* reconstituted complex confirmed that SNOR binds to the PTC of the 60S subunit at a site that is indistinguishable from its location in the *in situ* cryo-ET structure of the hibernating ribosome (Fig. 2d-f, Extended Data Fig. 5-6). Therefore, we first built and refined an atomic model of the SNOR-containing 60S complex using the high-resolution SPA cryo-EM map. We then applied rigid-body fitting of the cryo-EM model together with the model of the *S pombe* 40S subunit^24^ into the lower-resolution cryo-ET map to obtain a consensus model of the hibernating ribosome (see Material and Methods).

### SNOR interactions with the rRNA are important for ribosome binding

SNOR binds within the canyon formed by the ribosomal tRNA-binding sites, capping the entrance to the polypeptide exit tunnel (PET) (Fig. 3a-b, Extended Data Fig. 5). On the ribosome, SNOR interacts almost exclusively with rRNA via charged interaction mediated by residues K68, H96, and R97 (Fig. 3d-f). In particular, it is positioned behind helix 69 (H69), stabilizing the noncanonical conformation of this helix. This positioning of SNOR also explains the previously observed inactive ribosome population^24^, which prevents H69 from folding backward into a canonical conformation.

**Figure 3.**
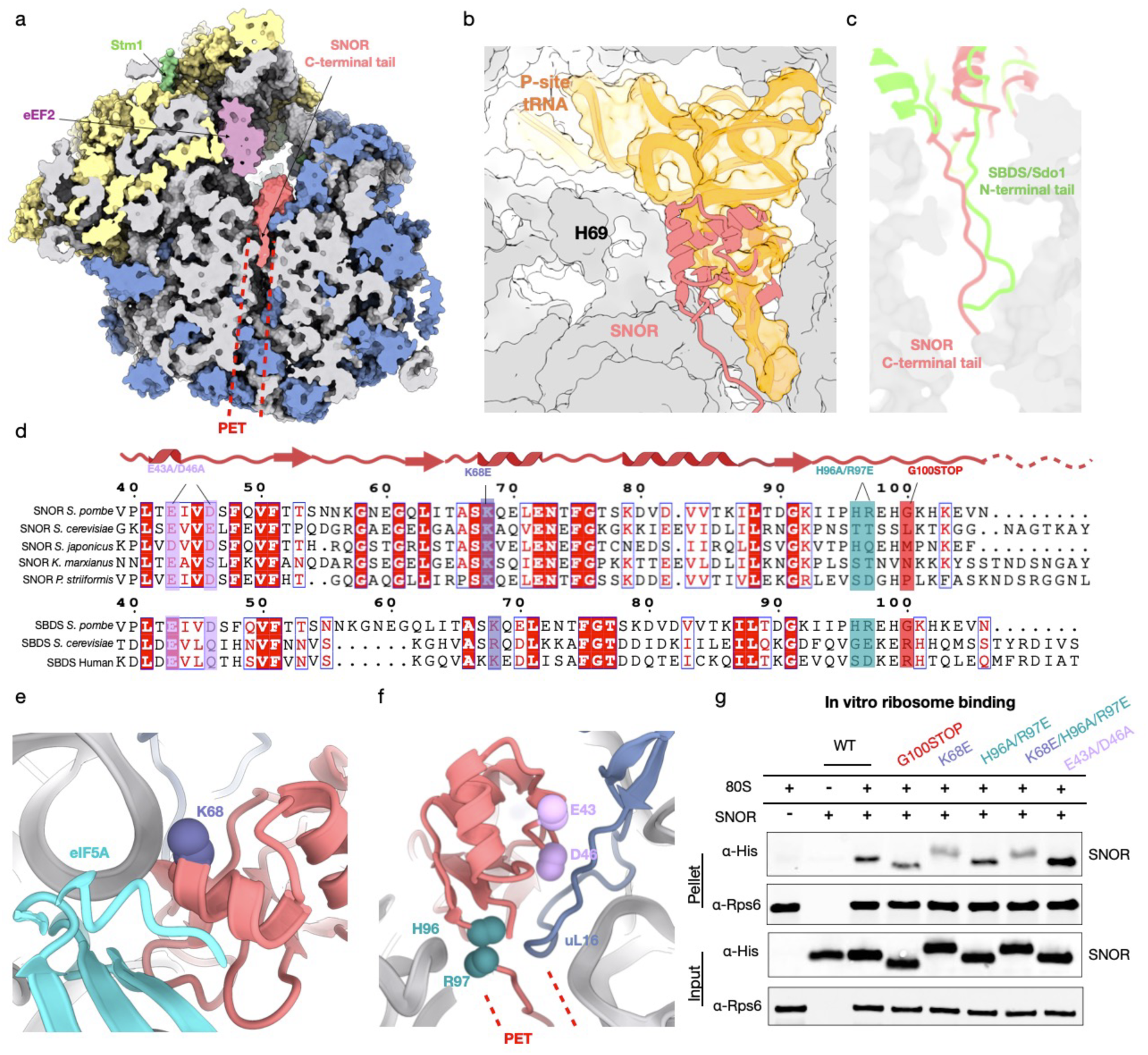
Structural and mutational analysis of SNOR-bound hibernating ribosome. (a) Slab view of the *S. pombe* hibernating ribosome atomic model shown as surface, highlighting insertion of the SNOR C-terminal tail into the polypeptide exit tunnel (PET, dashed line). Ribosomal RNA is shown in gray; small subunit proteins in yellow; large subunit proteins in blue; eEF2 in purple; Stm1 in green; SNOR in coral. (b) Structural comparison of SNOR and P-site tRNA binding site (PDB: 9AXV). Ribosomal protein and RNA are shown as a gray surface; SNOR is depicted as a coral cartoon; P-site tRNA is shown in orange. (c) Comparison of SNOR and SBDS tail (PDB: 6QKL) insertion into the PET. Atomic models of SNOR and SBDS are shown in coral and green, respectively. (d) Multiple sequence alignment of *S. pombe* SNOR with homologs from four fungal species (top) and with *S. cerevisiae* and human SBDS (bottom), highlighting conserved ribosome-contacting residues: E43 and E46 (lilac), K68 (purple), H96 and R97 (teal), and G100 (coral), which was selected for C-terminal truncation. Alignment generated using ESPript 3.0. (e) Close-up view of SNOR residue K68 (purple) contacting ribosomal RNA helix. (f) Close-up of SNOR residues E43 and E46 (lilac) contacting ribosomal protein uL16 (blue), and residues H96 and R97 (teal) contacting ribosomal RNA. PET location indicated by dashed red line. (g) Immunoblot analysis of ribosome co-sedimentation assay showing interaction of wildtype SNOR and SNOR mutants (G100STOP, K68E, H96A/R97E, K68E/H96A/R97E, and E43A/D46A) with purified *S. pombe* 80S ribosomes. SNOR was detected via His-tag; Rps6 served as a loading control.

In addition to its interactions with rRNA, SNOR establishes multiple contacts with the ribosomal protein uL16 (Rpl10) (Fig. 3f). Notably, uL16 contains a flexible loop that is disordered in the empty ribosome^31^ but adopts an ordered conformation when specific factors like tRNA or Sdo1 engage the ribosome^25,32^. This loop extends toward the P-site and interacts with the CCA end of the P-site tRNA during translation^32^. In the current structure, however, this loop establishes contacts with a loop located between alpha helix 2 and beta-sheet 3 of SNOR. Residues E43 and D46 of SNOR interact with uL16. This interaction is reminiscent of the one observed between the SBDS domain of Sdo1 and the immature 60S subunit^25^.

To assess the conservation of these critical residues contacting the ribosome, we performed a multiple sequence alignment of SNOR across various yeast species, also including Sdo1 and human SBDS proteins for further comparison (Fig. 3d). Most residues interacting with the ribosomal RNA and proteins were conserved in yeast. In particular, K68 showed strong conservation, including with human SBDS, which contains two lysine residues in this region, one of which has been reported as a disease-associated K/E variant^33^. To then evaluate the functional relevance of these interactions, we generated a series of SNOR mutants and then tested their binding to the ribosome. First, we probed SNOR interactions with the rRNA. We generated a single mutant (K68E), a double mutant (H96A/R97E), and a triple mutant (K68E/H96A/R97E). We assessed ribosome-binding activity using *in vitro* co-sedimentation assays followed by immunoblot analysis (Fig. 3g). The K68E single mutant and K68E/H96A/R97E triple mutant both exhibited reduced ribosome binding relative to wildtype SNOR, while the H96A/R97E double mutant demonstrated no change in its binding to the ribosome. These results suggest that highly conserved K68 is the primary contributor to RNA-mediated SNOR binding to the ribosome.

The C-terminal tail of SNOR, which is composed of seven residues enriched in positively charged amino acids, was observed inserting into the PET (Extended Data Fig. 7). This mode of interaction contrasts with that of Sdo1, which inserts its N-terminal domain into the PET^25^ (Fig. 3c and 2f, and Extended Data Fig. 7). To test the importance of this tail, we generated truncation mutant (G100STOP) lacking the C-terminal tail to test the role of PET insertion. The C-terminal truncation had also reduced ribosome binding, albeit to a lesser extent than K68E mutant. This indicates that PET insertion is not required for association but may serve another role, perhaps sensing nascent peptide presence within the tunnel. Finally, a double mutant (E43A/D46A) was created to disrupt interactions with uL16. Interestingly, this mutant displayed enhanced binding to the ribosome, possibly due to the removal of negatively charged residues that might otherwise repel the surrounding rRNA.

### SNOR is involved in translation repression

SNOR contacts eIF5A in the vicinity of the PTC, whereas eIF5A simultaneously engages with the ribosomal protein uL1, forming a tripartite interface (Fig. 4a-b). eIF5a engagement with uL1 stabilize the L1 stalk in an inward (closed) position (Fig. 4c), which is consistent with the previous structure of stalled ribosomes containing eIF5A^32^. Notably, the uL1–eIF5A–SNOR tripartite interface may act as a wedge that locks interactions between H69, the PTC, and the L1 stalk. This configuration could help maintain a hibernation-like state by preventing L1-tRNA interactions that normally occur during the elongation cycle^34,35^. Based on this interaction and the critical position of SNOR on the ribosome, we hypothesized that it plays a role in repression of protein synthesis in response to glucose depletion.

**Figure 4.**
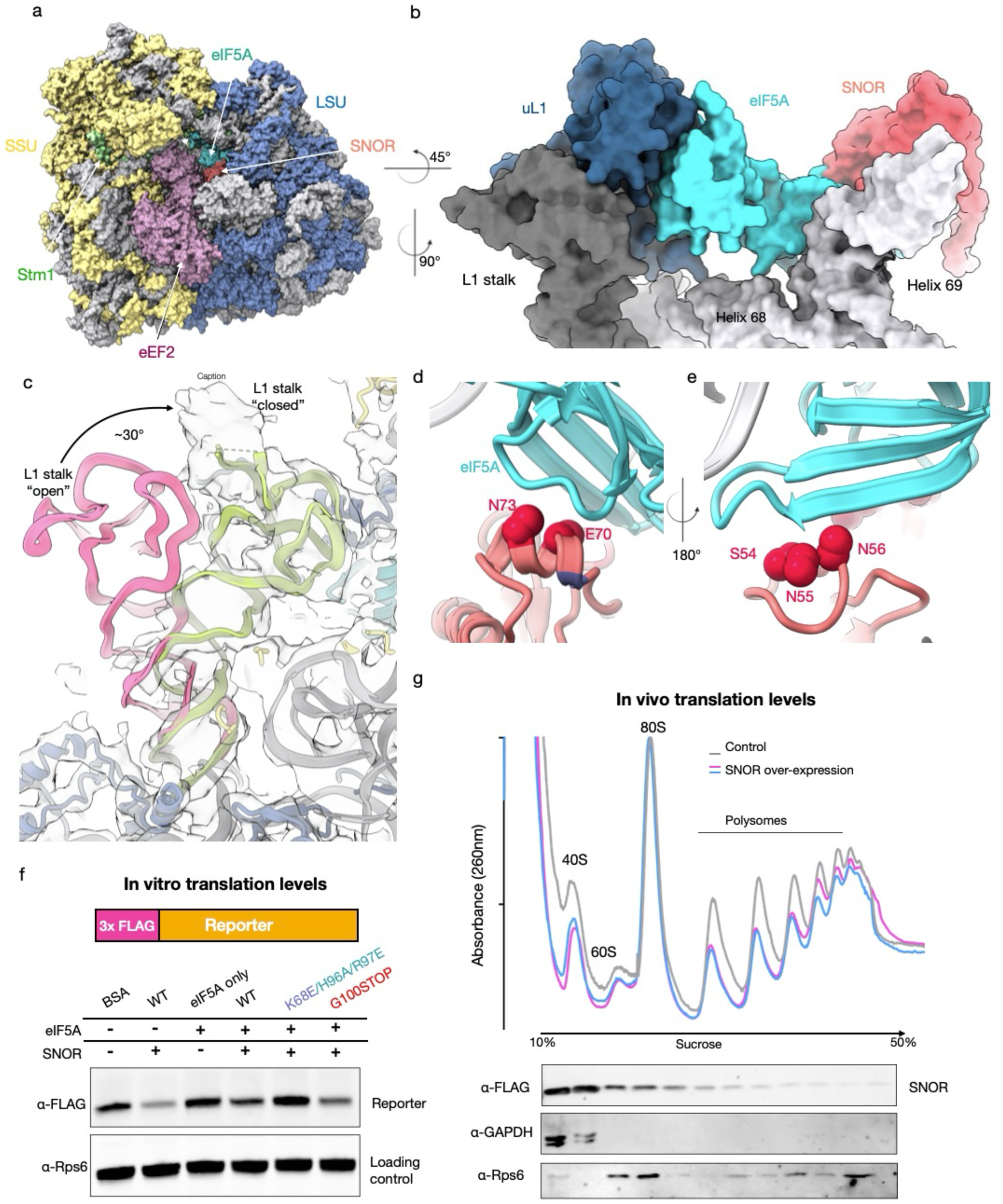
SNOR-eIF5a-uL1 tripartite interface stabilizes the ribosome L1 stalk and H69. (a) Atomic model of the *S. pombe* hibernating ribosome shown as a surface representation. Ribosomal RNA is depicted in gray; small subunit proteins in yellow; large subunit proteins in blue; eEF2 in purple; Stm1 in green; eIF5A in teal; SNOR in coral. (b) Close-up view of the SNOR–eIF5A–L1 tripartite interface locking the L1 stalk. SNOR is shown in coral; eIF5A in teal; ribosomal protein uL1 in navy; rRNA helices 68 and 69 in gray; the L1 stalk is highlighted in charcoal. (c) Conformational comparison of the L1 stalk in “open” (pink, PDB: 6WOO) versus “closed” (green) states. The ribosome atomic model is shown as cartoon; the cryo-ET map shown as a semi-transparent surface. (d-e) Close-up views of potential SNOR–eIF5A interaction interfaces involving SNOR residues S54/N55/N56 and E70/N73 (shown in red) and eIF5A (teal). (f) *In vitro* translation assay using rabbit reticulocyte lysate (RRL) comparing FLAG-tagged reporter levels in the presence of wildtype SNOR and mutants and/or eIF5A, compared to BSA control, assessed by immunoblot, done as independent duplicates. (g) Overlay of polysome gradient profiles from *S. pombe* cells grown in EMM with 0.5% glucose for 3 days under non-expressing (control, gray) and expressing (blue and pink) conditions, measured by absorbance at 260 nm (top). Distribution of SNOR, Rps6 and GAPDH across gradient fractions was assessed by immunoblotting (bottom).

To assess whether SNOR can inhibit translation, we used *in vitro* translation system using rabbit reticulocyte lysate (RRL) and monitored protein synthesis using a FLAG-tagged nascent chain construct. We confirmed that SNOR also associates with mammalian ribosomes using *in vitro* binding assays with purified human ribosomes from HEK cells, followed by co-sedimentation through a sucrose cushion (Extended Data Fig. 8). We then performed *in vitro* translation in RRL in the presence or absence of SNOR. The addition of purified SNOR to the RRL reaction significantly reduced FLAG-reporter levels compared to the BSA control (Fig. 4f). Next, we asked whether the addition of eIF5A would further enhance translation repression SNOR. The addition of eIF5A alone into RRL showed increased FLAG-reporter levels consistent with its role for increasing translation efficiency^32^, which was reduced by the addition of SNOR. In contrast, the SNOR triple mutant (K68E/H96A/R96E) that poorly binds ribosomes failed to repress translation, while G100 mutants showed a partial reduction in reporter synthesis, consistent with its weaker ribosome binding (Fig. 4f).

Next, we assessed whether SNOR represses translation *in vivo*. To this end, we overexpressed SNOR in *S. pombe* and monitored global translation by polysome profiling. SNOR expression led to a ∼30% decrease in polysome levels relative to 80S monosomes, indicating a modest translational slowdown compared to control cells (Fig. 4g). We also validated SNOR binding to ribosomes *in vivo* using immunoblotting of the ribosomal fractions from the sucrose gradients (Fig. 4g). Altogether, our results indicate that SNOR participates in translation repression under nutrient stress.

### SNOR restarts protein synthesis following cell dormancy

Since the impact of SNOR observed on translation repression is limited, we hypothesized that SNOR plays an additional role in restarting translation after the exit from cellular dormancy. Previously, we showed that S. pombe cells grown in EMM with low glucose (0.5% w/v) stop dividing and shut down protein synthesis after 3–4 days, once glucose is exhausted, as indicated by the loss of polysomes in sucrose gradients and transition to dormancy by day 7 ^24^.

To examine SNOR’s role in this process, we generated an *rtc3Δ* (*SNOR* KO) strain and compared its growth to wildtype under different glucose conditions (2%, 0.5%, and 0.02%). The knockout strain showed no major growth defect in any of these conditions (Fig. 5a). We then incubated both strains in 0.5% glucose for 7 days and assessed protein synthesis using polysome profiling. By day 7, polysomes had shifted into 80S monosomes, confirming global translation shutdown (Extended Data Fig. 9). When glucose was reintroduced by switching to glucose-containing media for 2 hours, polysomes reappeared in wildtype cells, indicating translation had restarted. However, *SNOR* KO cells failed to show polysome recovery, suggesting a defect in restarting protein synthesis (Fig. 5b and c; Extended Data Figs. 9 and 10).

**Figure 5.**
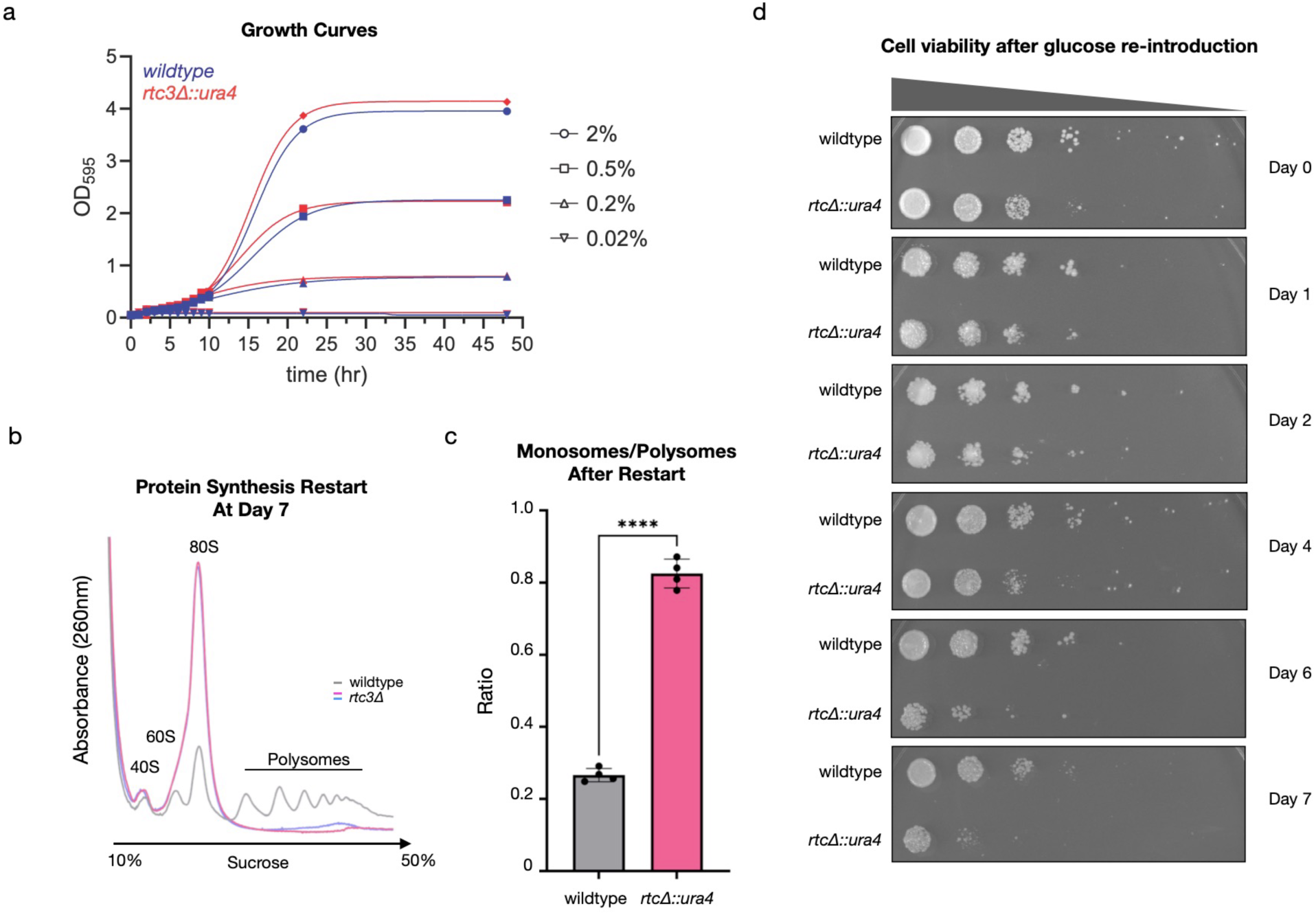
SNOR is essential for cell viability and for the restart of protein synthesis after glucose reintroduction. (a) Growth curves of *wildtype* and *rtc3Δ* cultures in EMM with different glucose levels at 32°C. (b) Overlay of representative polysome gradient profiles from *S. pombe* cells after reintroduction of glucose for 2 hours following 7 days of glucose depletion. Profile obtained from wildtype cells shown in gray, profiles obtained from ΔSNOR cells shown in pink and blue. Statistical analysis was performed using t-test.****P < 0.0001. (c) Quantification of relative area under the curve of 80S monosome levels (panel B). Data shown as mean ± S.D. (d) Serial dilution spot assays of cultures grown to saturation in EMM with 0.5% glucose at 32°C and then continuously incubated in the same media. 10-fold serial dilutions of the normalized samples at OD595 of 0.25 were spotted on EMM plates containing 2% glucose and incubated at 32°C. Day 1 corresponds to the 24 h time point of continuous incubation.

To further investigate the physiology of this defect, we grew wildtype and *SNOR* KO cells in low glucose media (0.5%) for up to 10 days and reintroduced them to fresh 2% glucose EMM plates to assess growth recovery. We collected samples every day, measured optical density (OD), and monitored viability by spotting cells on plates. For days 0-2, both wildtype and KO strains showed similar cell viability (Fig. 5d and Extended Data Fig. 11a). However, starting on day 4, *SNOR* KO cells began to show reduced viability compared to wildtype. Notably, this is consistent with the observed increase in SNOR protein levels at day 4 (Fig. 2b) and also with the onset of protein synthesis shutdown ^24^. By days 6 and 7, the knockout strain exhibited a significant viability defect relative to wildtype, and the *rtc3Δ* cells exhibited a more granulated and vacuolated morphology (Extended Data Fig. 11b). These findings are also consistent with the polysome gradient analysis and corroborated a failure to restart protein synthesis (Fig. 5b).

Combined, our results demonstrate that SNOR is essential for cell recovery from nutrient stress once glucose becomes available again and is required for exit from dormancy.

### SNOR expression is sensitive to glucose stress and is conserved across Fungi

To determine whether SNOR induction is specific to glucose stress or part of a broader response to nutrient imbalance, we subjected cells to a range of nutrient conditions, including limiting (0.5% w/v) and non-limiting (2% w/v) glucose concentrations. Since one of the natural habitats for S. pombe is honeybee honey^36^, which contains glucose levels varying between 20 to 30%, we also assessed SNOR expression under high glucose conditions (20% w/v). Finally, we tested complete glucose starvation (0%), which is known to rapidly suppress protein synthesis^37–39^, as well as amino acid and nitrogen starvation, which triggers distinct nutrient stress responses^40,41^, including a recently discovered 40S subunit degradation pathway ^42^.

Immunoblot analysis revealed that SNOR expression was selectively upregulated under low (0.5%) and also high glucose (20%) conditions but remained unchanged under normal glucose levels (Fig. 6a). Notably, SNOR levels did not increase under 0% glucose. This condition leads to rapid translational shutdown within minutes ^39^, therefore, we speculate this may likely prevent the induction and synthesis of SNOR. Similarly, SNOR expression was not affected by amino acid or nitrogen starvation, suggesting that its regulation is not part of a general response to nutrient limitation. Together, these results indicate that SNOR is specifically upregulated in response to stress under limiting or high glucose conditions but not to other nutrient stress signals such as nitrogen and amino acids.

**Figure 6.**
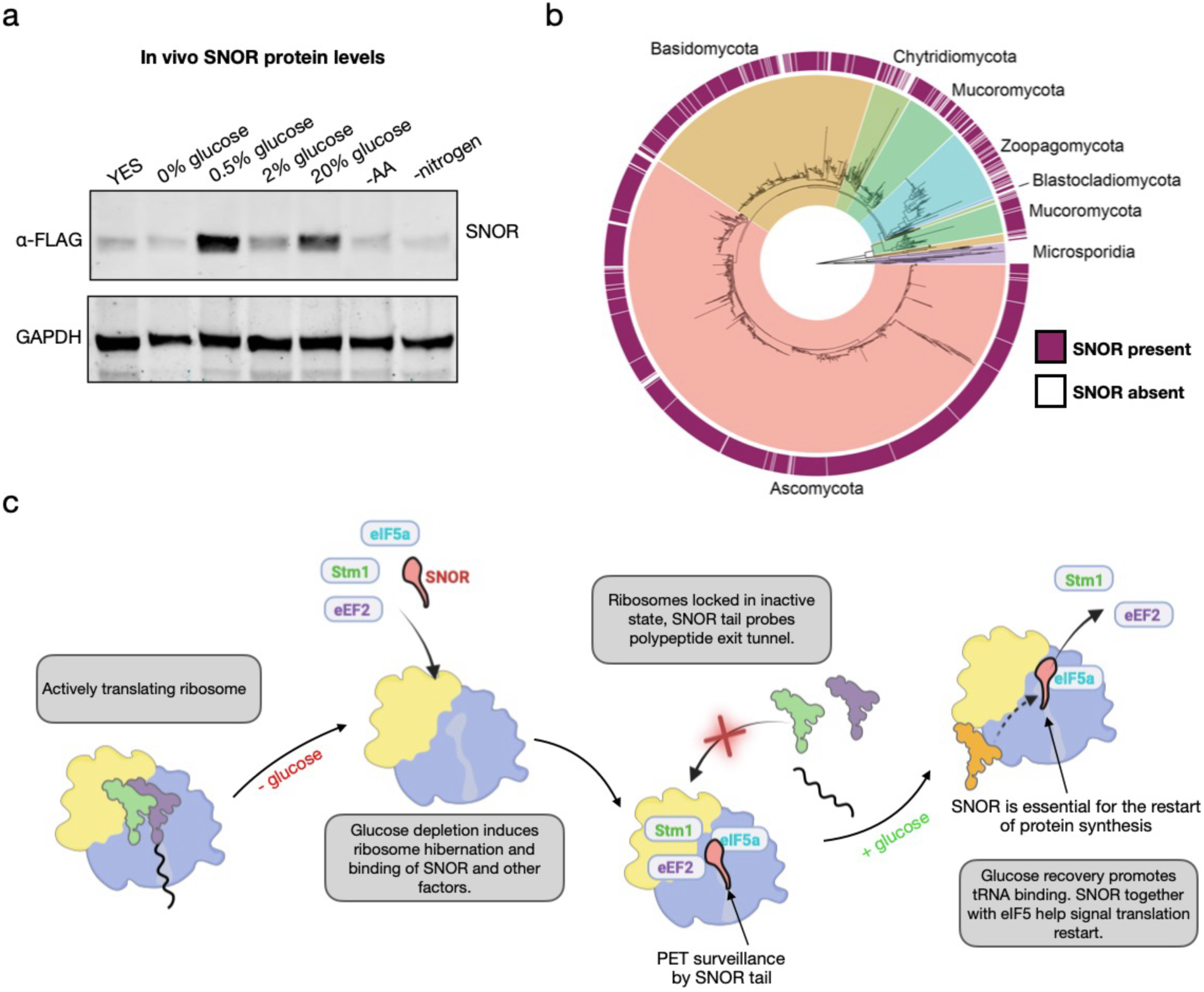
SNOR is a conserved factor specifically expressed in response to high and limiting glucose levels. (a) Immunoblot analysis of SNOR levels in *S. pombe* cell lysates exposed to various stress conditions for 2 hours before cell lysis. SNOR was detected using anti-FLAG antibody; GAPDH served as a loading control. (b) An 18S rRNA maximum likelihood of 2,248 fungal genomes showing the distribution of SNOR. The heatmap corresponds with the detection of SNOR within each genome. (c) Model for SNOR function in protein synthesis repression and restart during glucose depletion and repletion.

Next, we asked how conserved SNOR is across fungi. To assess its distribution across fungal species, we searched 2,248 representative fungal genomes from the NCBI RefSeq database using SNOR sequences highly similar to the protein characterized in this study. SNOR was detected in 87% of surveyed fungi and was predominantly found in Ascomycota, Basidiomycota, and Mucoromycota (94%, 87%, and 85% of annotated genomes, respectively) (Figure 6b, Extended Data Table 3 and 4). Most genomes surveyed encoded a single SNOR homolog, though a few contained up to five copies. Conversely, SNOR was entirely absent in Microsporidia, which are known for their highly compact genomes resulting from extreme genome reduction associated with obligate intracellular parasitism^43^.

Initially, we did not detect clear SNOR homologs in plants, animals, or other multicellular eukaryotes, suggesting that SNOR may be a fungi-specific factor. To assess whether SNOR homologs exist in other eukaryotes, we expanded the survey to humans and other mammal genomes. SNOR was undetected in the annotated proteins of 263 mammalian genomes we searched. Interestingly, we found several hits in plants, insects, fish and in some mammals that were annotated as an SBDS N-terminal domain containing protein. However, they were only detected at transcript level and contain low sequence homology to SNOR (<30%), therefore we excluded these hits from our final search criteria. Together, our data indicates that SNOR overexpression is a response to glucose stress and is highly conserved in fungi.

## Discussion

Cells faced with environmental stress often switch to dormancy as a survival strategy to persist under harsh environmental conditions. Energy-intensive processes such as protein synthesis are selectively shut down to preserve resources until favorable conditions are restored. In this study, we leveraged cellular cryo-ET and SPA cryo-EM, together with complementary genetic and biochemical analyses to visualize hibernating ribosomes in their native cellular environment and discovered a 12-kDa stress-induced factor, SNOR.

Our results demonstrate that after glucose depletion, all hibernating ribosomes become associated with SNOR, which, in concert with factors such as Stm1 (Oga1), eIF5A, and eEF2, plays a central role in translational shutdown (Fig. 6c; Extended Data Movie 1). Our data indicate that SNOR can repress translation by occupying the PTC and entrance to the PET, while simultaneously bridging interactions between these factors. Importantly, once glucose levels are restored, SNOR, potentially with the sequestration of eIF5A on the ribosome, signal the restart of protein synthesis. This model helps explain why multiple hibernation factors are required not only to coordinate ribosome inactivation, but also for protein synthesis reactivation and confer specificity to distinct metabolic states, rather than relying on a single dominant regulator.

Our structural data reveal that SNOR shares homology with Sdo1 (SBDS in humans), a ribosome biogenesis factor during late stages of assembly^33,44,45^. Intriguingly, SNOR appears to repurpose its SBDS domain not for ribosome maturation, but to probe the PTC environment in mature ribosomes. In addition, SNOR inserts its C-terminal tail into the ribosomal exit tunnel. By occupying this space, SNOR can sense potential nascent chains stuck in the tunnel of hibernating ribosome prior to restart of translation. A recently identified hibernation factor Dap1b formed of unstructured tail was found bound to the PET of hibernating ribosomes isolated from unfertilized eggs in Zebrafish and Xenopus^21^. Unlike SNOR, Dap1b lacks a structured domain outside the tunnel (Extended Data Fig. 12), which suggests it may perform a different function.

Another unexpected feature of SNOR is its interaction with eIF5A. eIF5A is a translation factor that contains a hypusine modification that is known to interact with tRNA to promote translation elongation and promote peptide-bond formation during various ribosome stalling events, including between consecutive proline residues^32,46,47^. A similar interaction was also observed between eIF5A and Dap1b (Extended Data Fig. 12) ^21^. In the current structure, however, SNOR and eIF5A together with the ribosomal protein L1 form a tripartite interface that bridges the L1 stalk to helix H69 of the rRNA. This interaction may help retain eIF5A on the ribosome which may be required to restart protein synthesis once glucose levels are restored. These findings expand the known roles of eIF5A beyond elongation, termination, and quality control to include a potential function in translational shutdown and restart during nutrient-induced stress.

Functionally, we determined that SNOR levels are upregulated in response to both low and high glucose levels, but not to other starvation conditions such as amino acids and nitrogen. Therefore, SNOR expression and its ribosome-binding activity support a model in which SNOR links glucose metabolism status to translational control. Finally, phylogenetic analysis shows that SNOR is highly conserved across fungi. This conservation highlights the biological importance of this protein in fungal species. Conversely, SNOR was almost entirely undetected in Microsporidia (Extended Data Table 3). Microsporidia have particularly small genomes, as they have undergone significant genome reduction to become obligate intracellular parasites^43,48^. The microsporidia-specific hibernation factor MDF1^49^, although structurally very different from SNOR, occupies a similar position and thus may serve an analogous role. Protein sequence search resulted in multiple results that show SBDS-containing protein paralogs exist in different eukaryotes, including plants, insects, and mammals. It remains to be determined if SNOR-like proteins are present in humans.

Our recent findings revealed that hibernating ribosomes can tether to mitochondria, forming organized lattices on the mitochondrial membrane^24^. While the functional significance of this tethering remains to be fully understood, it is possible that SNOR acts in response to change in mitochondria bioenergetics and glycolysis to coordinate global translational shutdown in the cytoplasm and on the mitochondria during cellular stress.

Our study identifies SNOR as a previously unrecognized, conserved regulator of translation on the eukaryotic ribosome that responds specifically to glucose stress and is essential for reactivating protein synthesis during recovery from cellular dormancy. SNOR provides an intriguing framework for understanding how cells modulate protein synthesis during the entry and exit from dormancy by linking metabolic state to surveillance performed on the ribosomal active site. This discovery also highlights the power of integrative structural biology and visual proteomics in uncovering unexpected layers of protein translational control.

## Materials and Methods

### Sample preparation for cryo-ET

Cryo-ET samples were prepared using 200-mesh R2/2 copper grids (Quantifoil) that were plasma-cleaned for 30 s in a 75% argon / 25% oxygen atmosphere with a 1070 plasma cleaner (Fischione). *Schizosaccharomyces pombe* cells were plunge-frozen using a Leica EM GP2 plunger, operated at 23 °C and 100% relative humidity. *S. pombe* cells were grown for 7 days in EMM containing 0.5% (w/v) glucose. Prior to freezing, cultures were diluted in glucose-free EMM to an OD600 of 0.6. A 4 µL aliquot of the diluted suspension was deposited onto the carbon side of the plasma-cleaned grid. Back-side blotting was performed for 1 s from the copper side using Whatman 597 paper, employing the paper contact sensing mode with an added movement of 1.5 mm. Grids were then vitrified in liquid ethane.

### Cryo-FIB milling for lamella preparation

Vitrified grids were mounted into CryoFIB Autogrids (ThermoScientific, ref. 1205101) under liquid nitrogen and transferred into an Aquilos 2 cryo-FIB/SEM system (ThermoScientific), operated via xT software. The temperature of the stage and the anti-contamination shield was maintained at −194 °C throughout the procedure. Grid screening was performed in the SEM mode at 5 kV and 13 pA. Grids were then coated for 120 s with an organometallic platinum precursor trimethyl(methylcyclopentadienyl)platinum(IV) using the integrated gas injection system (GIS), preceded and followed by a 15 s sputter coating with metallic platinum at 30 mA. Lamellae were prepared using AutoTEM for automated milling, with the ion beam set at 30 kV and progressively reduced currents from 1 nA to 30 pA. Final thinning to 160 nm was performed manually at 30 pA. During the last milling steps, the stage was tilted by an additional +0.5° to enable top surface polishing of each lamella, reducing thickness irregularities and minimizing curtaining artifacts. A final coating layer of platinum was applied by sputtering (3s at 10 mA) to reduce sample charging during TEM acquisition and help with TS alignment^50^.

### Cryo-ET data acquisition in glucose-depleted wildtype cells

Data collection was carried out on a Titan Krios G4i transmission electron microscope (ThermoScientific) operating at 300 kV, equipped with an energy filter and a Falcon4i direct electron detector used in zero-loss mode. Initial mapping of lamellae was done at a pixel size of 14.39 Å, using a defocus of −70 µm, a 70 µm objective aperture, and a 20 eV energy slit. Mitochondria were identified for tilt series (TS) acquisition in SerialEM^51^, guided by lamella segmentation and selection/optimisation using SPACEtomo^52^. Reference images were further acquired to refine beam image-shift alignments for parallel acquisition using PACEtomo^53^. Tilt series were collected at a nominal magnification of 64,000×, corresponding to a calibrated pixel size of 1.933 Å on the detector. A 50 µm C2 and a 70 µm objective aperture were used, with the energy filter slit narrowed to 10 eV. The microscope was operated in nanoprobe mode with spot size 5 and an illumination area of 1.4 μm. TS were acquired following a dose-symmetric scheme with 3° tilt increments, spanning from −65° to +49°, centered relative to the lamella pre-tilt (approximately −8°). Each movie was recorded in counting mode over 880 ms, with an average dose of ∼3.34 e^−^/Å² per movie, with raw fractions saved in eer format. The total electron dose per tilt series was ∼130 e^−^/Å². Target defocus values ranged from −2 µm to −6 µm, with 0.5 µm increments between series. A total of 78 tilt series were selected for subtomogram averaging (STA), based on criteria including optimal dose rate, minimal lamella thickness, lack of contamination, and absence of signs of cellular apoptosis.

### Cryo-ET data processing

The workflows for cryo-ET data processing and subtomogram averaging are outlined in Extended Data Figure 1. As a first step, stage tilt angles recorded in the .mdoc metadata files were adjusted to account for lamella pre-tilt, ensuring that reconstructed tomograms were flat and that the tilt series were centered around the tilt angle corresponding to the highest electron dose (minimal thickness at 0). Raw movie frames were converted from .eer to .tiff format using the relion_convert_to_tiff tool in RELION v4.0.1^54^. Motion correction and frame averaging were then performed, followed by contrast transfer function (CTF) estimation using a defocus range of −2 to −10 µm in Warp 1.09^55^. Images and their associated CTF fits were inspected manually, and poor-quality projections were excluded. Tilt series stacks were assembled and aligned using AreTomo2^56^. Finally, defocus handedness was verified before reconstructing 3D-CTF corrected tomograms at a final pixel size of 15.44 Å using Warp.

### Ribosome picking and selection of OMM-tethered ribosomes

Ribosomes were identified in the 3D-CTF corrected tomograms using template matching implemented in PyTOM^57^. As a reference, a low-pass filtered (40 Å) EM map of an *S. pombe* outer mitochondrial membrane (OMM)-associated ribosome (EMD-50266) was used. To distinguish OMM-tethered ribosomes from free cytosolic ones, spatial masks dilated around the mitochondrial membranes were employed to filter the template-matching hits. For generating these masks, tomograms were first reconstructed in Warp with deconvolution. Semantic segmentation of two tomograms was performed in Dragonfly (Comet Technologies), and the resulting annotations were used to train a 2.5D U-Net (five-slice input, depth level 5, patch size 64 pixels) to segment mitochondrial membranes and adjacent ribosomes. This trained model was subsequently applied in batch mode to predict membranes and generate dilation masks that encompassed tethered ribosomes. These masks were applied during particle extraction to isolate OMM-associated ribosomes from the full set of template-matched particles. The extracted particles were then manually curated in ArtiaX^58^ to remove false positives and unbound cytosolic ribosomes.

### Subtomogram averaging, classification, and multi-particle refinement

Subtomograms of the selected OMM-tethered ribosomes, along with their corresponding CTF information, were extracted in Warp and used to compute an initial average, which served as the starting reference for 3D refinement in *RELION*. Additional rounds of 3D classification without alignment and subsequent refinement steps were carried out to discard low-quality particles and enhance alignment accuracy. The refined particle set was then processed in M^59^ for joint optimization of particle poses, local sample deformations, and CTF parameters. To further refine the dataset, cryoDRGN-ET^60^ was employed for unsupervised particle classification, allowing a finer classification of structurally heterogeneous particles. The resulting classes were used to generate different species in *M* for additional rounds of refinement. The final structure of the OMM-associated ribosome achieved a global resolution of 6.0 Å, based on the Fourier shell correlation (FSC = 0.143) criterion. Similar rounds of refinements and classification were performed for the unbound cytosolic ribosomes (identified by removing the tethered ribosomes from the template matching picks and further visual curation in ArtiaX), reaching a resolution of 6.6 Å. Once we confirmed the presence of the same hibernation factors bound in both tethered and free cytosolic ribosomes (Extended Data Fig. 2), we combined, refined and averaged the datasets in M, reaching a global resolution of 5.5 Å.

### Real-time qPCR

*S. pombe* cells were grown overnight at 30 °C with agitation in YES (Yeast Extract with Supplements) medium, then transferred to EMM (Edinburgh Minimal Medium) supplemented with 0.5% (w/v) glucose and cultured for an additional 3 days. Culture samples were harvested daily and flash frozen in liquid nitrogen for subsequent analysis. RNA was extracted from 1 x 10^7^ cells per sample, using the RNeasy Kit (QIAGEN, Cat. no. 74104) following the manufacturer’s protocol. Quantity and quality of RNA samples were assessed using a NanoDrop (Thermo Scientific). cDNA was synthesized using the Verso cDNA Synthesis Kit (Thermo Scientific, Cat. no. AB1453A). Real-time qPCR was performed on the StepOne Plus system (Applied Biosystems) using SYBR Green Master mix (Thermo Scientific, Cat. no. A25742). Gene expression levels were quantified using the ΔΔCt method and normalized to the expression of the housekeeping gene, *β-actin*. Primer sequences are listed in Extended Data Table 2.

### Monitoring SNOR protein levels during nutrient stress

*S. pombe* cells were grown overnight at 30 °C with agitation in EMM (Edinburgh Minimal Medium) medium. The overnight cultures were washed three times with PBS (Phosphate Buffered Saline) and used to inoculate fresh EMM (Edinburgh Minimal Medium) supplemented with either 0; 0,5 or 2% glucose or EMM lacking either amino acids or nitrogen but supplemented with 2% glucose. Cells were inoculated to a starting OD600 of 0.1 and cultured at 30°C with constant agitation for 40 minutes after which samples were collected for cell lysis. For cell lysis 1.5 OD600 units of cells were resuspended in 500μL of water. 50μL of 1.85 M NaOH was added and incubated on ice for 10 minutes. Trichloroacetic acid (TCA) was then added to a final concentration of 10%, followed by an additional 10-minute incubation on ice. Samples were centrifuged at 14,000 × g for 15 minutes at 4 °C. Pellets were then resuspended in 1X SDS-PAGE sample buffer (50 mM Tris-HCl pH 6.8, 2% SDS, 1% β-mercaptoethanol, 6% glycerol, 0.004% bromophenol blue) for Western blot analysis. Proteins were resolved using SurePAGE gels (GenScript, cat. no. M00653) with MES running buffer (GenScript, cat. no. M00677) and transferred onto 0.2 μm nitrocellulose membranes (LI-COR, cat. no. 926-31092). Membranes were stained with Ponceau S solution for 5 minutes, rinsed with distilled water, and imaged to assess protein loading. Afterward, membranes were blocked for 1 hour at room temperature in 5% milk prepared in 1X PBST (0.1% Tween-20). After blocking, antibodies diluted in 2% milk in 1X PBST were incubated with membranes. Primary antibodies used: FLAG (Genscript, cat no. A00187), GAPDH (Proteintech, cat no. 60004-1-Ig). Secondary antibodies used: anti-mouse (Invitrogen, cat no. A21058). The LI-COR Odyssey imager was used for detection.

### Ribosome purification

*S. pombe* cells were cultured overnight at 30 °C with agitation in 500 mL of YES (Yeast Extract with Supplements) medium. Cells were harvested by centrifugation at 3,000 × g for 5 minutes at room temperature, washed with Lysis Buffer (20 mM HEPES pH 7.4, 100 mM KCl, 5 mM MgCl₂, 1% (v/v) Triton X), and centrifuged again at 5,000 × g for 5 minutes. The resulting pellet was resuspended in Lysis Buffer supplemented with and 0.04 U/µL RNase inhibitor (RiboLock, Thermo Fisher, cat. no. EO0382) and flash frozen in liquid nitrogen as small drops. Frozen pellets were ground to a fine powder in a mortar and pestle under liquid nitrogen. The powder was resuspended 1:1 (v/w) in Lysis Buffer and the lysate was clarified by centrifugation at 5,000 × g for 5 minutes at 4 °C to remove cell debris, followed by a second centrifugation at 14,000 × g for 10 minutes at 4 °C. The resulting supernatant was layered over a 50% sucrose cushion (50% w/v sucrose, 20 mM HEPES pH 7.4, 100 mM KCl, 5 mM MgCl₂) and centrifuged at 43,000 rpm (50.2 Ti rotor, ∼225,000 × g) for 20 hours at 4 °C. Following centrifugation, the ribosome pellet was resuspended in either Resuspension Buffer (20 mM HEPES pH 7.4, 60 mM KCl, 5 mM MgCl₂) or Ribosome Splitting Buffer (20 mM HEPES pH 7.4, 1 M KCl, 5 mM MgCl₂, 1 mM puromycin). For subunit separation, ribosome samples were incubated in Ribosome Splitting Buffer for 1 hour at 4 °C, then loaded onto a continuous 10–40% sucrose gradient (10–40% w/v sucrose, 20 mM HEPES pH 7.4, 100 mM KCl, 5 mM MgCl₂, 1 mM DTT) and centrifuged at 21,000 rpm for 20 hours at 4 °C. Gradients were analyzed using a BIOCOMP Piston Gradient Fractionator™ system, and fractions corresponding to the small and large ribosomal subunits were collected. Subunit samples were concentrated using Amicon® Ultra Centrifugal Filter Concentrator tubes (100 kDa MWCO, Millipore, ref. UFC8100).

### Recombinant SNOR purification

A pET28a vector encoding N-terminal 6xHis-tagged SNOR was expressed in *E. coli* BL21-CodonPlus (DE3) competent cells (Agilent, cat. no. 230245). Site-directed point mutations in SNOR were generated using the QuickChange mutagenesis kit (Agilent, cat. no. 210518) and validated by whole plasmid sequencing by Plasmidsaurus. Cells were cultured in LB medium (Fisher Bioreagents, cat. no. BP1426) at 37 °C until reaching an OD600 = ∼0.6. Protein expression was induced with 1 mM IPTG, followed by overnight incubation at 18 °C. Cells were harvested by centrifugation, and the resulting pellet was resuspended in Lysis buffer (50 mM HEPES pH 7.5, 1 M KCl, 10% glycerol, 6 mM β-mercaptoethanol, 0.5x protease inhibitor cocktail) and lysed using a French Press. The lysate was clarified by two rounds of centrifugation at 20,000 rpm for 30 minutes at 4 °C using a TI 50.2 rotor. The cleared supernatant was loaded onto a 5 mL HisTrap HP column (Cytiva, cat. no. 17524802) using a P1 pump at 4 °C and washed with 1–2 column volumes (CV) of Wash Buffer (50 mM HEPES-KOH pH 7.5, 500 mM KCl, 15 mM imidazole, 10% glycerol, 6 mM β-mercaptoethanol). The column was then transferred to an ÄKTA Pure fast protein liquid chromatography system (Cytiva) and washed with an additional 5 CV of Wash Buffer. Bound proteins were eluted using a step gradient with 2–3 CV of 15%, 30% and 80% Elution Buffer (50 mM HEPES-KOH pH 7.5, 500 mM KCl, 300 mM imidazole, 10% glycerol, 6 mM β-mercaptoethanol). Fractions corresponding to the 30% and 80% elution peaks were pooled and transferred to 7 kDa MWCO dialysis tubing (ThermoFisher, cat. no. 27968700), then dialyzed overnight at 4 °C in dialysis buffer (50 mM HEPES-KOH pH 7.5, 150 mM KCl, 30 mM imidazole, 10% glycerol). The protein samples were subsequently concentrated using Amicon® Ultra Centrifugal Filter Concentrator tubes (3 kDa MWCO, Millipore, ref. UFC8003), aliquoted, flash frozen in liquid nitrogen, and stored at –80 °C.

### Ribosome co-pelleting assay

The 40S, 60S and 80S ribosome samples were thawed on ice and diluted to 1µM using Binding Buffer (50mM HEPES pH 7.7, 150 mM KOAc, 15 mM Mg(OAc)2, 0.02% C12E8, 0.04 U/µL, 5% v/v glycerol). Purified wildtype SNOR and SNOR point mutants were thawed on ice and diluted to 75 µM using the Binding buffer. To prepare the reactions, the purified ribosome samples were added to a final concentration of 0.2µM and the purified SNOR samples were added to a final concentration of 2µM. The reactions were incubated for 30 minutes at 30°C and layered 1:1 over a 40% sucrose cushion (40% w/v sucrose, 20 mM Hepes pH 7.4, 100 mM KCl, 5 mM MgCl2). Samples were centrifuged at 100,000 rpm for 2 hours at 4°C. Following the spin, the supernatant was discarded and the resulting pellet was resuspended in 1X SDS-PAGE sample buffer (50 mM Tris-HCl pH 6.8, 2% SDS, 1% β-mercaptoethanol, 6% glycerol, 0.004% bromophenol blue) for analysis by Western Blot. Samples were resolved on a SurePAGE gel (Genscript, cat no. M00653) using MES buffer (Genscript, cat no. M00677) and transferred onto a 0.2 μm nitrocellulose membrane (LI-COR, cat no. 926-31092). Membrane was blocked in 5% milk in 1X PBST (0.1% Tween-20) for 1hr at room temperature. After blocking, antibodies diluted in 2% milk in 1X PBST were incubated with membranes. Primary antibodies used: 6xHis (Genscript, cat no. A00186), RPS6 (Cell Signaling, cat. no. 2217). Secondary antibodies used: anti-mouse (Invitrogen, cat no. A21058), anti-rabbit (Invitrogen, cat no. A32735). The LI-COR Odyssey imager was used for detection.

### Cryo-EM sample preparation and data collection

The large ribosomal subunit and recombinant SNOR protein were purified as described above. Freshly prepared 60S ribosome samples were diluted to 270 ng/µL in resuspension buffer A (20 mM HEPES KOH pH 7.4, 60 mM KCl, 5 mM MgCl2). To reconstitute the 60S:SNOR complex, purified SNOR protein was first initially diluted using buffer A and then added to the reaction to a final concentration of 1 µM. The reaction was incubated for 30 minutes at 30°C and then moved to ice. Quantifoil Cu200 R2/2 grids were coated with a 3 nm carbon layer using a Safematic carbon coater (Rave Scientific) prior to sample application. Grids were plasma cleaned for 15 s at 15 mA using a Pelco easiGlow™ system to render the surface hydrophilic. Five µL of the reaction mixture were then applied to the grids, followed by a 60 s incubation. Grids were blotted for 9 seconds at blot force +7 and plunge-frozen in liquid ethane using a Vitrobot Mark IV (ThermoScientific) operated at 4 °C and 100% humidity. Cryo-EM data acquisition was carried out using a Titan Krios transmission electron microscope (ThermoScientific) operated at 300 kV. The microscope was equipped with a K3 direct electron detector and a Gatan Quantum energy filter set to a 10 eV slit width. Movies were recorded in counting mode at a nominal magnification of 105,000×, yielding a calibrated pixel size of 0.83 Å. A total of 7,277 movies were acquired, each consisting of 40 frames and accumulating a total electron dose of 50 e⁻/Å². The target defocus range was set between -1.6 and -0.6 μm.

### Cryo-EM data processing

Cryo-EM data processing was performed as outlined in Extended Data Fig. 4, using CryoSPARC for all steps. Movies were subjected to motion correction, dose-weighting, and contrast transfer function (CTF) estimation. Micrographs were curated to exclude damaged or low-quality exposures, resulting in 6,457 micrographs used for further analysis. Blob picker was used to select particles within a diameter range of 150Å and 400Å. Following inspection, a total of 849,708 particles were extracted from the curated micrographs using a box size of 512 × 512 pixels. These particles were subjected to 2D classification into 150 classes, from which 27 high-quality classes comprising 502,382 particles were selected. The selected particles underwent two rounds of heterogeneous refinement to remove junk particles, yielding a subset of 211,256 particles. These were used for non-uniform refinement, producing an initial reconstruction of the large ribosomal subunit and SNOR complex at a global resolution of 2.71 Å. Subsequently, 3D Variability Analysis (3DVA) was performed using a spherical mask around SNOR to identify particles with the most stable density for this region. A refined subset of 53,130 particles was selected and subjected to an additional round of non-uniform refinement, resulting in a final map with a global resolution of 2.91 Å. Resolution was determined by gold-standard Fourier Shell Correlation (FSC) between two independently refined half-sets, using a 0.143 threshold.

### Model building

#### Large ribosomal subunit and SNOR complex

Following data processing, a previously generated model of the *S. pombe* large ribosomal subunit (PDB: 9AXU) and AlphaFold prediction model of SNOR (AF-Q9P7K6-F1-v4) were docked into the cryo-EM map using ChimeraX. SNOR was further rigid body fit into the observed density using COOT and manually adjusted based on the observed density. PHENIX was used to refine the model in the 60S:SNOR map with 5 macrocycles of real space refinements applying Ramachandran, sidechain rotamer, protein secondary structure and nucleotide restraints to correct for clashes. The final model was validated using MolProbity in PHENIX. All figure generation was done using ChimeraX.

#### Hibernating Schizosaccharomyces pombe 80S ribosome

The previously obtained model of the *S. pombe* ribosome (9AXV) was docked into the *in situ* cryo-ET map using ChimeraX. As the fit of the AlphaFold predictions for *S. pombe* eEF2, eIF5A, and Stm1 into the observed density was not satisfactory, we used the PHYRE2 one-to-one threading option^61^ to align the *S. pombe* sequences of these factors to the structures of their respective *S. cerevisiae* homologs. The resulting homology models provided improved fits to the observed density and were subsequently used for model building. The resulting models of *S. pombe* eEF2, eIF5a and Stm1 along with the model of SNOR obtained from cryo-EM analysis of the large ribosome subunit - SNOR complex, were docked into the cryo-ET density using ChimeraX. COOT was used to further fit the observed factors as rigid bodies based on the densities observed in the map. The region of the L1 stalk was manually adjusted in COOT based on the observed density. The atomic model of the hibernating *S. pombe* 80S ribosome was refined against the *in situ* cryo-ET density map using 5 macrocycles of real-space refinement in PHENIX applying Ramachandran, sidechain rotamer, protein secondary structure and nucleotide restraints to correct for clashes. The final model was validated using MolProbity as implemented in PHENIX.

### Small-scale in vitro translation

Purified mRNA of N-terminal 3x Flag-tagged reporter was translated in rabbit reticulocyte lysate (RRL; Promega, cat. no. L4540). The lysate was diluted to 66.7% (v/v) with a translation mix containing purified RNA (final concentration 0.5 µg/µL), 3 µM SNOR, 3 µM eIF5A, a combination of both, or 3 µM BSA as a control. The final reaction buffer also included 0.04 U/µL RNase inhibitor (Promega), 0.5× protease inhibitor cocktail (Promega), 81 mM KCl, 2 mM magnesium acetate, and 24 µM amino acid mix. Reactions were incubated at 32 °C for 25 minutes, then moved on ice. Following the incubation, samples were analyzed by SDS-PAGE and western blotting to detect levels of FLAG-tagged translation product.

### In vivo experiments

#### SNOR KO and mutants strain generation

To construct the *rtc3Δ::ura4^+^* strain, 300 bp of the *rtc3^+^* 5’ UTR and 3’UTR were amplified from the genomic DNA of wildtype cells. The resulting amplicons were cloned into the BamHI/PstI site and KpnI/XhoI site, respectively, of a pSK plasmid containing the *ura4^+^* gene within the PstI/KpnI sites using the Gibson assembly method. A PCR product containing the UTRs and *ura4^+^* was then transformed into *ura4-D18* cells using a lithium acetate method^62^. Transformants were selected on Edinburgh minimal media (EMM) agar plates lacking uracil and the correct deletion was verified by whole-cell PCR using oligonucleotides flanking the *rtc3* UTR sequences and internal to *ura4^+^*.

Differential interference contrast images were acquired using a Zeiss Axio Observer inverted epifluorescence microscope with Zeiss 63× oil (1.46 NA) objective and captured using Zeiss ZEN 3.0 (Blue edition) software. A singular medial Z slice was obtained. All images were further processed using ImageJ (Schindelin et al., 2012). PMC3855844

#### Genomic FLAG tagging of SNOR

To add two FLAG tags to the N-terminus of SNOR, a CRISPR-Cas9-based approach was used. A guide RNA (gRNA) targeting a region near the SNOR start codon was selected using https://crispr.dbcls.jp/ and cloned into a Cas9 gRNA plasmid to induce a double-stranded break at the desired genomic location. The plasmid backbone was a generous gift from Dr. Henry Levin (NIH). Cloning was performed using Q5 polymerase, following the manufacturer’s instructions. In parallel, a double-stranded repair template was synthesised as a gBlock, containing in order: 200 nucleotides of the SNOR 5′ untranslated region (5′UTR), two FLAG tags beginning with AUG, and 200 nucleotides of the SNOR coding sequence (CDS). *S. pombe* cells (*S. pombe* YHL 912 h-, ura4-294 leu1-32 gift from Dr. Henry Levin) were then co-transformed with the gRNA-Cas9 plasmid and the repair template. The transformation protocol was adapted from Levin et al.^63^. After transformation, cells were plated on a selective medium (EMM) lacking leucine and incubated for 3 days. Next, individual colonies were transferred to YES medium to allow for the loss of the gRNA-Cas9 plasmid. Colonies were checked for FLAG tag incorporation by colony PCR as well as Western Blot.

#### Overexpression of SNOR

To generate yeast strains overexpressing SNOR, a plasmid placing N-terminal 2x-FLAG-tagged *rtc3^+^* under the control of the nmt1 promoter was used^64^. The pREP3 plasmid backbone was kindly provided by Dr. Henry Levin (NIH). Cloning was performed using Q5 polymerase and the Gibson Assembly method, following the manufacturer’s instructions. The pREP3X(pHL1766) plasmid was used as a backbone and was kindly provided by Dr. Henry Levin (NIH). Cloning was performed using Q5 polymerase and the Gibson Assembly method, following the manufacturer’s instructions. *S. pombe* strain YHL 912 *h^-^, ura4-294 leu1-32* was used for the overexpression experiments. Transformation was carried out using the protocol adapted from Levin et al^63^, and transformants were selected on selective medium (EMM) lacking leucine. To induce SNOR overexpression, cells were cultured in a selective medium (EMM) lacking thiamine. As a negative control (non-expressing condition), cells were grown in medium supplemented with thiamine at a final concentration of 5 µg/mL. Cells were grown in EMM supplemented with low glucose (0.5% w/v) for 3 days under either non-expressing (with 5 µg/mL thiamine) or expressing (without thiamine) conditions. After 3 days, cells were harvested for polysome gradient profiling (see below).

#### Glucose restart experiment and polysome gradient profiling

Wildtype and *rtc3Δ* mutant cells were grown to mid-log phase at 32°C in EMM containing 2% glucose overnight, adjusted to the same OD595 of 0.05 in 0.5% glucose EMM, (considered day 0 of the experiment) and then incubated continuously for the remainder of the experiment at 32°C. Each day, an aliquot was removed from each culture and adjusted to an OD595 of 0.25. By 3 days when cell proliferation had stopped, the same culture dilutions were made as at day 3. 10-fold serial dilutions of the normalized samples were spotted on EMM plates containing 0.5% glucose and incubated at 32°C. A technical replicate was performed at each time point. All samples were spotted on the same plate, and plates were imaged 3 days later. The experiment was performed twice.

For analyzing polysome gradients, cells were first incubated in EMM supplemented with 0.5% (w/v) glucose for 7 days. Following glucose depletion, cells were harvested, washed with double distilled H2O, resuspended in fresh YES (Yeast Extract with Supplements) medium to an OD600 of 1 and incubated for 2 hours in 30°C with agitation. Following the incubation, cells were treated with 100 μg/mL cycloheximide for 15 minutes, then harvested and lysed as described above (see Ribosome Purification). The clarified lysate was quantified by measuring absorbance at 260 nm, and 10 A260 units were loaded onto a continuous 10–50% sucrose gradient. Gradients were centrifuged at 230,000 × g for 2.5 hours at 4 °C. Polysome gradient profiles were analyzed using a BIOCOMP Piston Gradient Fractionator™ and visualized with GraphPad Prism 10.

### Gene detection and phylogenetics

Annotated protein sequences for all fungal (2,248 genomes) and mammalian (263 genomes) reference genomes (available on GitHub at https://github.com/cassprince/SNOR_conservation) were downloaded from the NCBI RefSeq database on December 28, 2024, and April 3, 2025, respectively. This resulted in a subset of RefSeq that includes approximately one genome per species^65,66^. SNOR protein sequences were detected using HMMER v3.3 (hmmsearch) (hmmer.org- Extended Data Table 4) with an E-value cutoff of 0.05. To search for SNOR, HMMER profiles were built with 6 protein sequences annotated as Rtc3 from Ascomycota genomes by the NCBI Eukaryotic Genome Annotation Pipeline^66^. Query sequences are given in Table S4. Only protein hits with lengths fewer than 200 amino acids were considered to be SNOR to minimize the likelihood of detecting larger aberrant SBDS proteins.

18S rRNA sequences from the fungal genomes described above were identified using BLAST v2.16.0^67^, aligned using MAFFT v7.520^68^, and applied to FastTree v2.1.10^69^ to produce a maximum-likelihood tree. The constructed tree was midpoint rooted with the phangorn v2.12.1 package^70^ and visualized with ggtree v3.12.0^71^. Taxonomic classifications were assigned according to the NCBI Taxonomy database^72^. The complete list of genome accession numbers, taxonomic classification, and SNOR presence for all surveyed genomes can be found in Table S5 and S6.

### 3D rendering of tomograms and EM volumes

Models of mitochondrial membranes displayed in Figure 1 and in the video (Extended Data Video 1) were segmented in Dragonfly and then imported into Blender 4.2 (https://www.blender.org/) using Microscopy Nodes plugin^73^, together with the corresponding tomogram. Ribosomes were placed in the 3D rendering layout following their positions in the .star files using the plugin Molecular Nodes^74^. All other EM volumes were prepared with ChimeraX.

## Acknowledgements

We are grateful to members of the Jomaa and Mattei labs for their valuable discussions. We thank Dr. Michael Purdy for his assistance with single-particle EM data collection and Dr. Alaina H. Willet for DNA primer design. Cryo-EM data collection was carried out at the Molecular Electron Microscopy Core (MEMC) at the University of Virginia School of Medicine. We also acknowledge the Imaging Centre at the European Molecular Biology Laboratory (EMBL IC) for access and support, made possible by generous funding from the Boehringer Ingelheim Foundation. We are thankful to the EMBL central IT services and to Thomas Hoffmann for their help with computational resources. This research was supported by The National Science Foundation Grant (Division of Molecular and Cellular Biosciences, Award # 2503218), by the Searle Scholars Program (Grant # SSP-2023-106), by The Owens Family Foundation, and aided by Grant #134088-IRG-19-143-33-IRG from the American Cancer Society awarded to A.J. Support for S.M. was provided by EMBL and to K.L.G by NIH grant R35GM131799. M.G. received funding through a Boehringer Ingelheim Fonds predoctoral fellowship, and H.R. was supported by the EMBL International PhD Program.

## Author Contributions

A.J., S.M., M.G., and H.R. conceived the study, M.G. purified the ribosomes and performed the biochemical experiments. H.R. performed cryo-FIB milling and cryo-ET data collection and processing. M.G. and M.P. collected and processed the cryo-EM data. M.B. performed gene tagging and overexpression in *S. pombe*. L.A.T. and K.L.G. generated *S. pombe rtc3* deletion and viability assays. C.R.P. and H.A.F. performed phylogeny analysis. A.J., M.G., S.M., and H.R. wrote the manuscript. All authors contributed to data analysis and the final version of the manuscript.

## Data Availability

Cryo-EM maps and model coordinates are deposited in the EMDB as EMD-XXXX and in the PDB as PDB ID YYY for the in-situ consensus cryo-ET hibernating ribosome, and EMD-XXXX and EMD-XXXX for the cryo-ET mitochondria-tethered and free cytosolic free hibernating ribosome, respectively, and EMD-XXXX and in the PDB as PDB ID YYY for the cryo-EM structure of the *S. pombe* 60S:SNOR ribosomes. All other data are available in the main text or the supplementary materials.

## Competing Interests

The authors declare that they have no competing interests.

## Notes

### Competing Interest Statement

The authors have declared no competing interest.

