## Extended Data and Tables for "A Novel Eukaryotic Ribosome Factor Enables Translation Restart Following Cellular Dormancy"

Gluc, Rosa *et al.*

Extended Data Figure 1-12

Extended Data Table 1-5

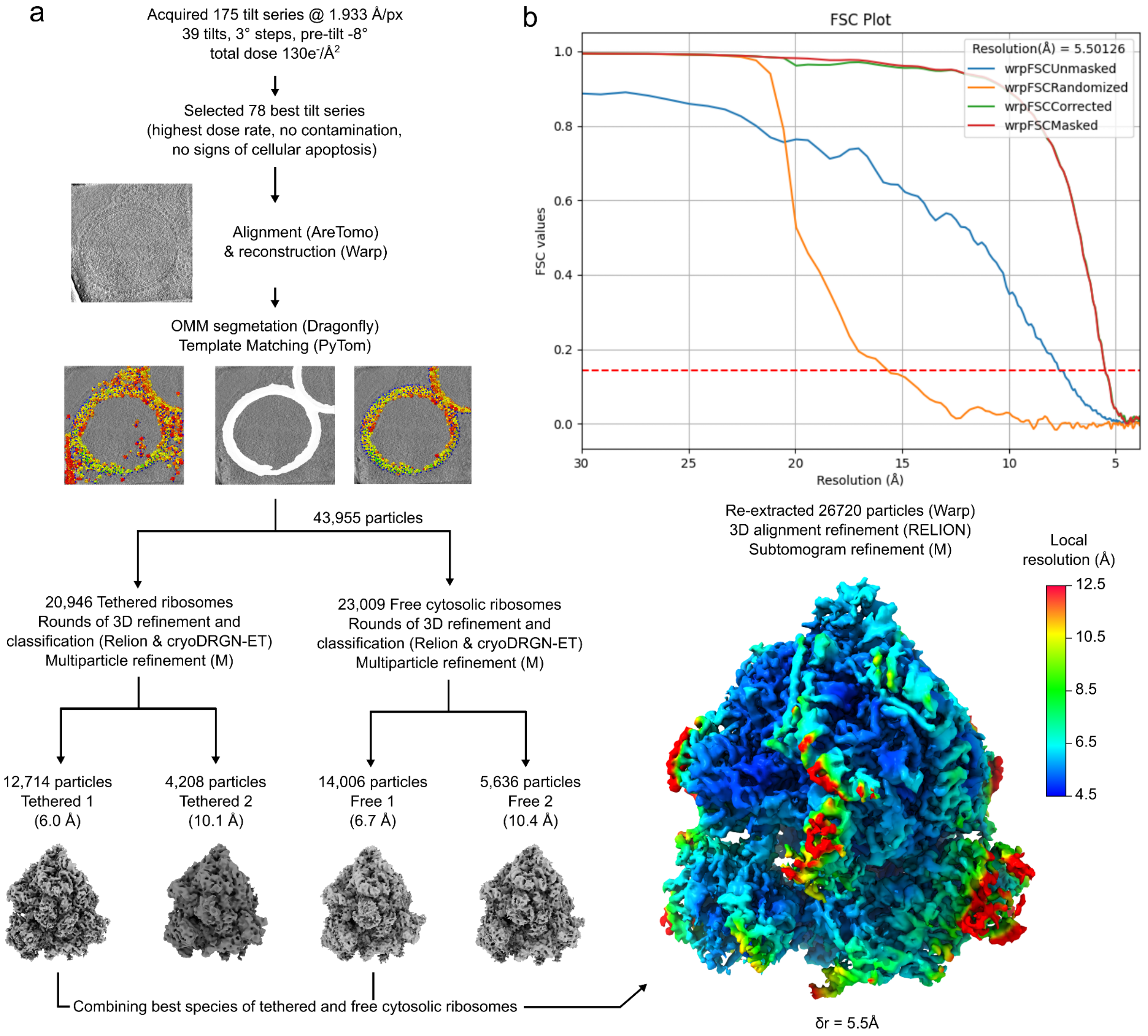


**Extended Data Figure 1. Processing scheme of *in situ* cryo-ET data.**

(a) Flow-chart of the cryo-ET pipeline used on WT *S. pombe* cells: following cell freezing at day 7 of glucose depletion and lamellae milling, dose-symmetric tilt series were acquired using a Titan Krios TEM. The 78 best tilt-series were aligned, reconstructed as 3D-CTF corrected tomograms (Using Warp). Segmentation (using Dragonfly) and template matching using (PyTom) allowed for the selection of OMM-tethered and free cytosolic ribosomes. After subtomogram extraction, consecutive rounds of 3D refinement and classification in RELION & cryoDRGN-ET, followed by multiparticle refinement in M. Finally, after carefully checking the presence of the same factors in the best classes for OMM-tethered and free cytosolic ribosomes, particles were re-extracted and combined for another round of 3D refinement in RELION and M, obtaining a density map of hibernation ribosomes with a global resolution of 5.5 Å, using 26,720 particles. (b) Fourier shell correlation (FSC) curve of the cryo-ET reconstruction of the *S. pombe* hibernating ribosome refined using M.

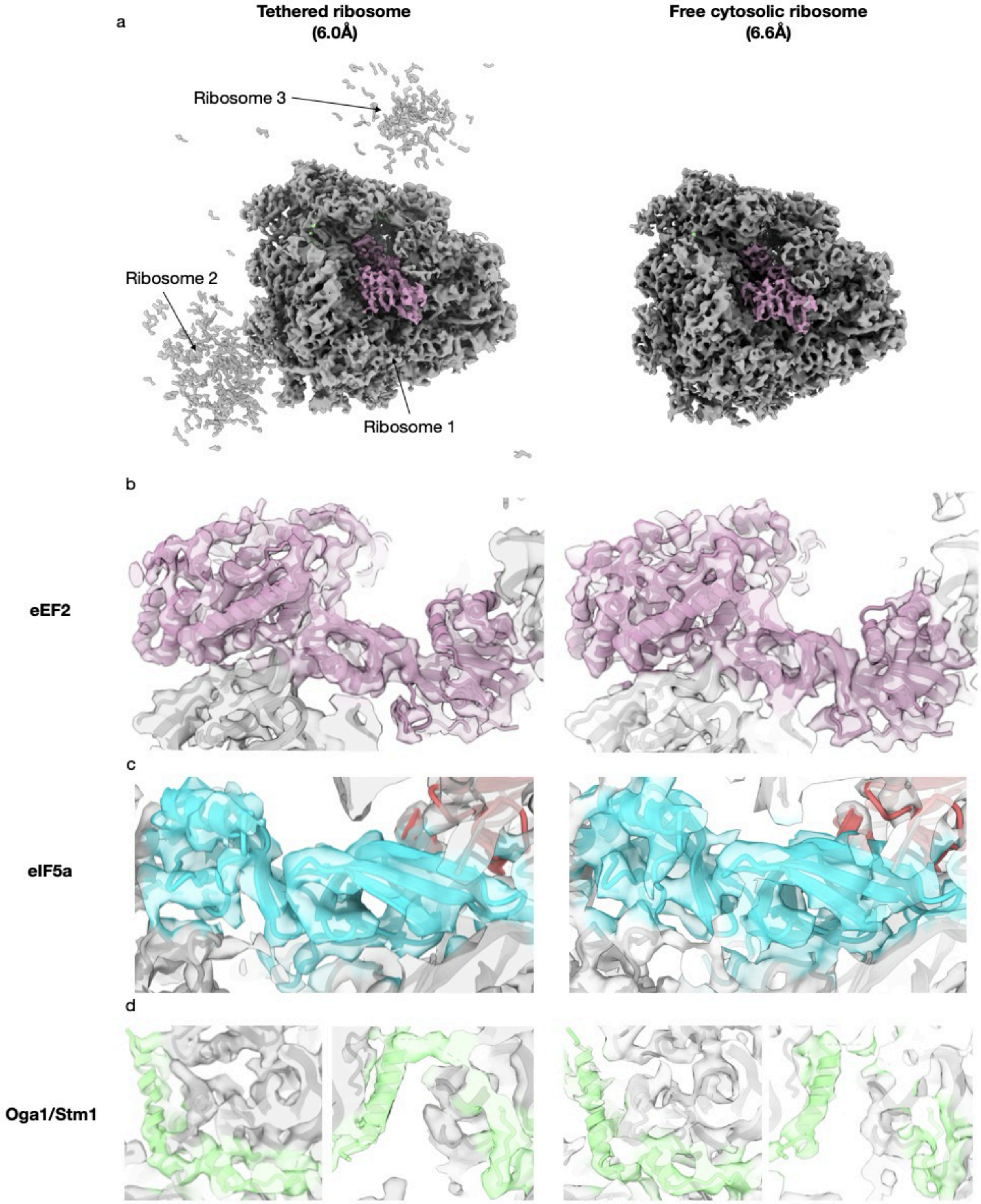
a
Tethered ribosome
(6.0Å)
Free cytosolic ribosome
(6.6Å)
Ribosome 3
Ribosome 2
Ribosome 1
b
eEF2
c
eIF5a
d
Oga1/Stm1

**Extended Data Figure 2. Densities of hibernation factors bound to the ribosomes in the OMM-tethered and free cytosolic ribosome maps.**

(a) Reconstructions of the *S. pombe* hibernating ribosomes obtained by subtomogram averaging of OMM-tethered (left) and free cytosolic (right) ribosome particles. Maps shown as surface, densities corresponding to observed hibernation factors shown in color. (b-d) Close-ups of densities corresponding to the observed hibernation factors bound to OMM-tethered ribosomes (left) and free cytosolic ribosomes (right). eEF2 shown in purple, eIF5a shown in teal, Oga1/Stm1 shown in green. Atomic models shown as cartoon, density maps shown as surface.

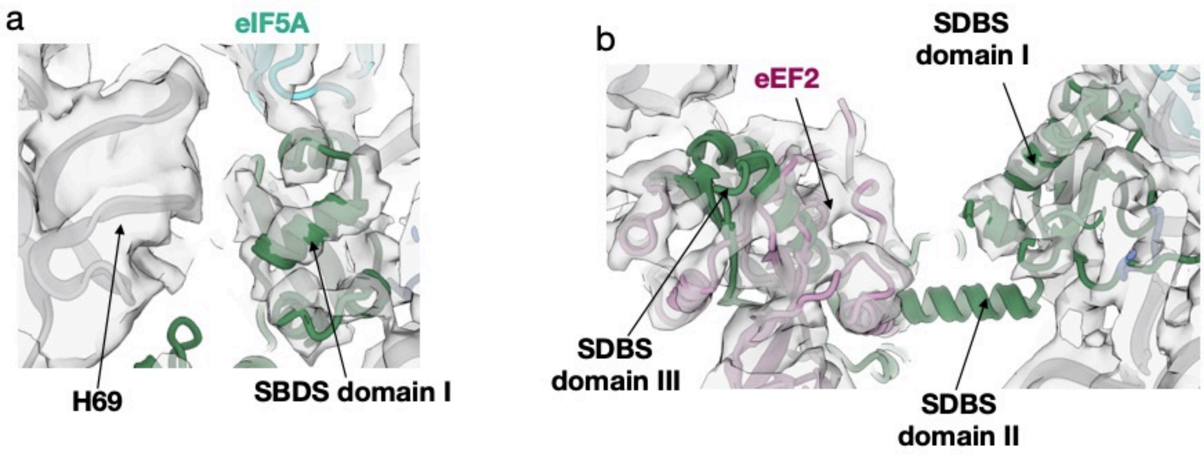


**Extended Data Figure 3. Domains I but not Domain II or III of Sdo1/SBDS fit into the EM-density of the hibernating ribosomes.**

(a) Fit of SBDS domain I (green) into the cryo-EM density of the *S. pombe* hibernating ribosome. Ribosomal RNA is shown in gray, with helix 69 indicated by an arrow; eIF5A is shown in teal. The electron density map is displayed as a semi-transparent surface. (b) Absence of observable density for SBDS domain II and predicted steric clash between SBDS domain III and eEF2 (purple). The positions of SBDS domains II and III are indicated by arrows. The electron density map is shown as a semi-transparent surface.

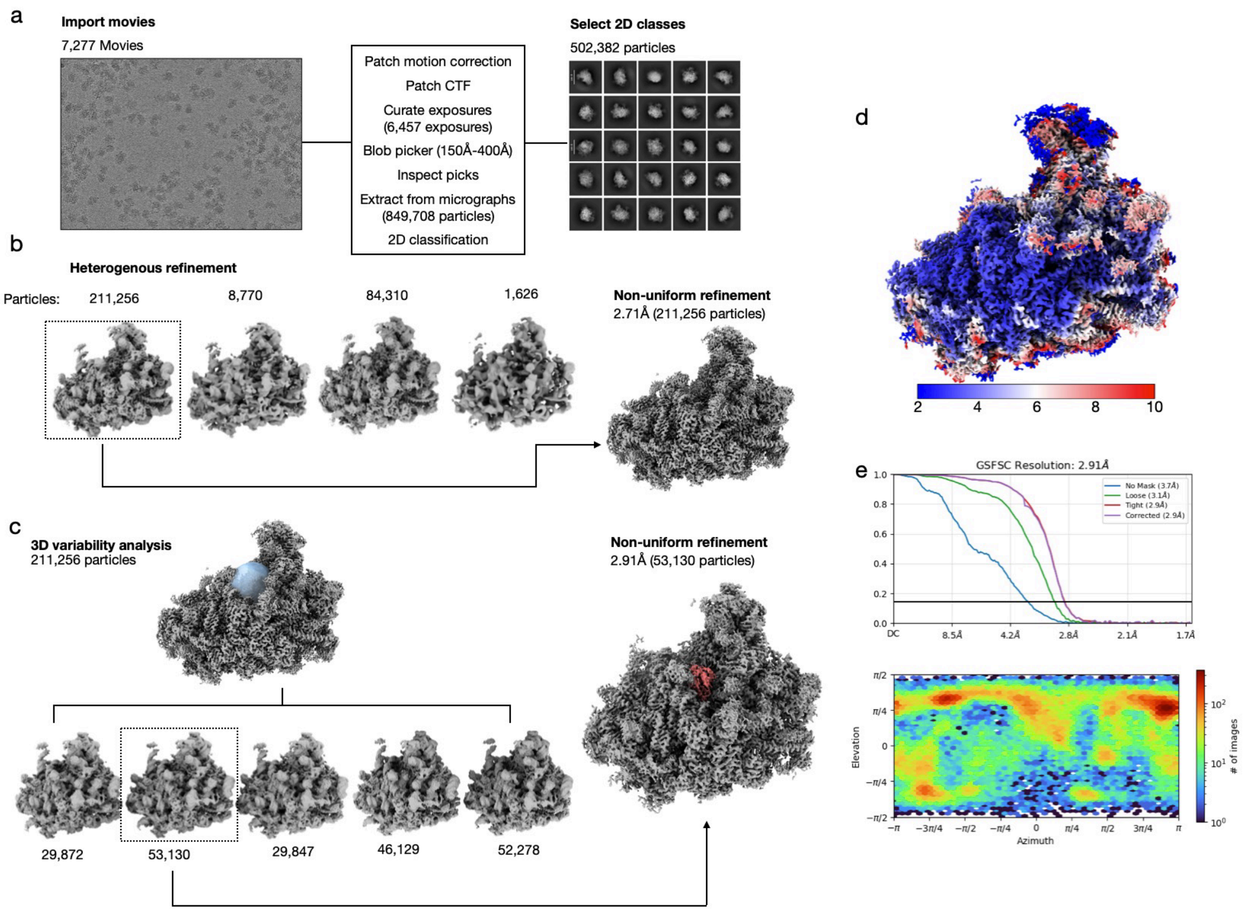


**Extended Data Figure 4. Processing scheme of the single-particle analysis of 60S:SNOR complex.**

(a) Motion corrected and dose-weighted micrographs were curated to remove damaged exposures. Blob picker was used to select for particles resulting in 849,708 initial picks. (b) Following 2D classification 502,382 particles were selected for heterogenous refinement yielding a class consisting of 211,256 used for further non-uniform refinement. (c) The consensus ribosome particles were subjected to 3D variability analysis using focused mask for SNOR (blue). Class consisting of 53,130 particles displaying most stable SNOR density was selected for non-uniform refinement resulting in a reconstruction of the 60S:SNOR complex at an average resolution of 2.91Å determined by gold standard Fourier Shell Correlation (GSFSC) at an FSC cutoff of 0.143. (d) 3D representation of local resolution calculated for the final 60S:SNOR complex map. (e)

Fourier Shell Correlation (FSC) plot for final map (top) and angular distribution plot (bottom) were generated in cryoSPARC.

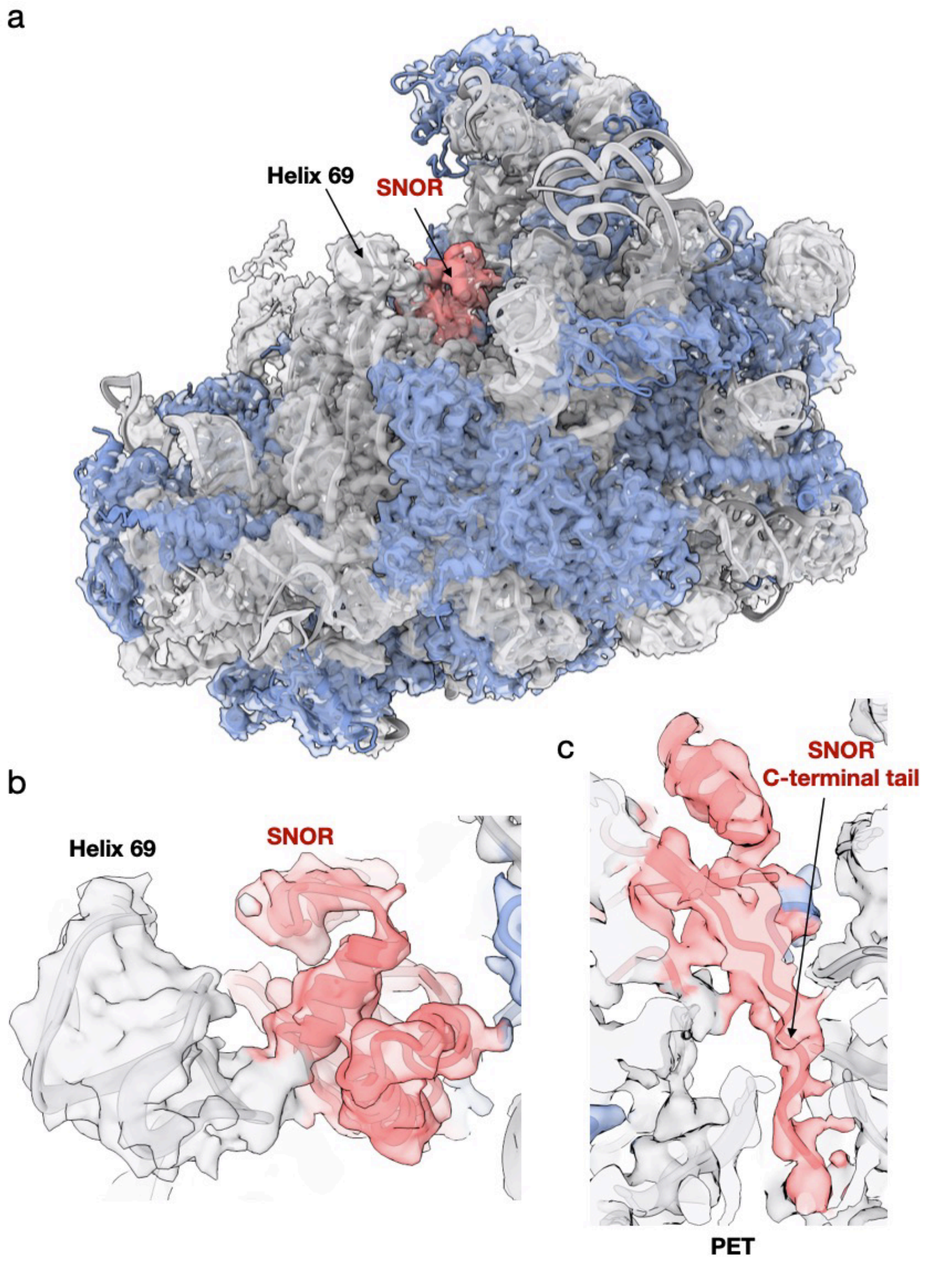


**Extended Data Figure 5. Atomic model of SNOR bound to the ribosomal large subunit.**

(a) Overall fit of the 60S:SNOR atomic model into the cryo-EM density map, shown as a semi-transparent surface filtered to 3.5 Å. Ribosomal RNA is shown in gray, ribosomal proteins in blue, and SNOR in coral. RNA helix 69 and the SNOR binding site are indicated by arrows. (b) Top-down close-up view showing the fit of SNOR within the observed density at the peptidyl transferase center (PTC), adjacent to helix 69. (c) Close-up view of the SNOR C-terminal tail inserted into the polypeptide exit tunnel (PET).

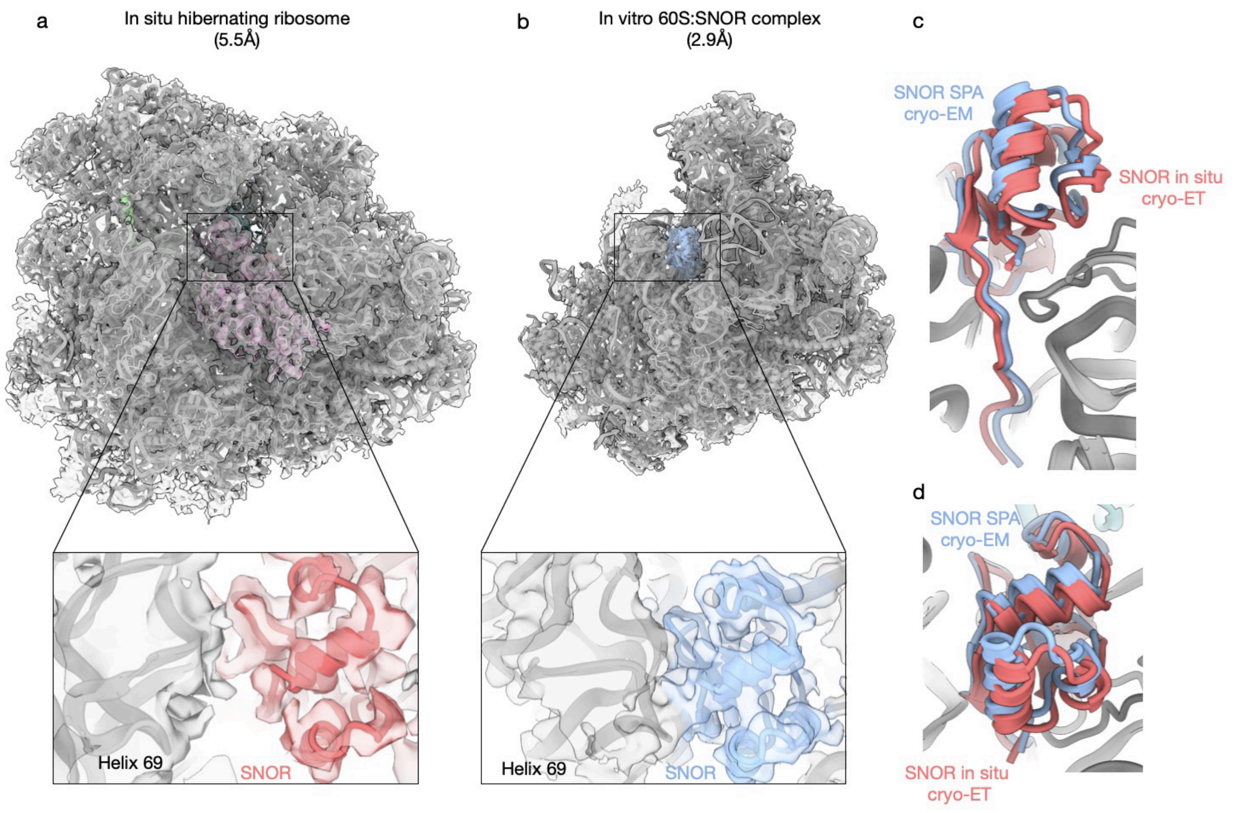


**Extended Data Figure 6. Comparison of SNOR binding position between *in situ* hibernating ribosome and *in vitro* reconstituted 60S:SNOR complex atomic models.**

(a) Overall fit of the atomic model of the *S. pombe* hibernating ribosome bound to eEF2 (purple), eIF5a (teal), Oga1 (Stm1, green) and SNOR (coral). Inset shows close-up of SNOR binding position in the PTC near helix 69 of the large ribosomal subunit RNA, density corresponding to SNOR colored coral. (b) Overall fit of the atomic model of *in vitro* reconstituted complex of the large ribosomal subunit and SNOR (sky blue). Inset shows close-up of SNOR binding position in the PTC near helix 69 of the large ribosomal subunit RNA, density corresponding to SNOR colored sky blue. (c-d) Overlay of atomic models of SNOR obtained from the *in situ* cryo-ET (SNOR colored coral) and *in vitro* SPA cryo-EM (SNOR colored sky blue) reconstructions. In all renderings the atomic models are shown as cartoon and the maps shown as semi-transparent surface. All images were produced in ChimeraX.

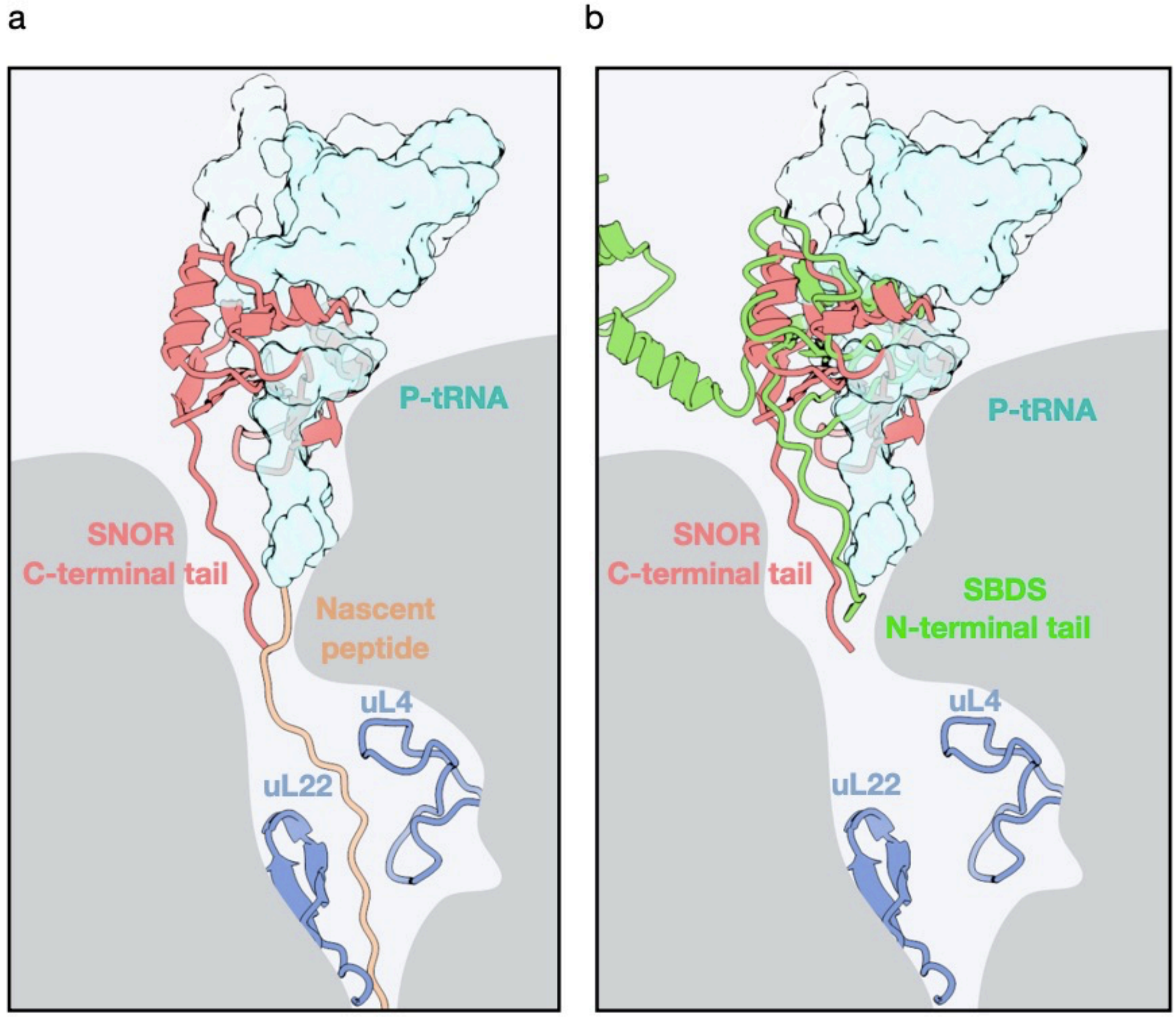


**Extended Data Figure 7. Schematic view of SNOR binding in the PET valley.**

(a) SNOR occupies the site of the P-site tRNA, with its C-terminal tail extending into the path of the nascent peptide (orange). (b) SNOR binding position overlaps with that of SBDS domain I (PDB: 6QKL); however, SBDS inserts its N-terminal tail into the PET (green). The P-site tRNA is shown as a light blue surface, and ribosomal proteins uL22 and uL4 are depicted as blue cartoons.

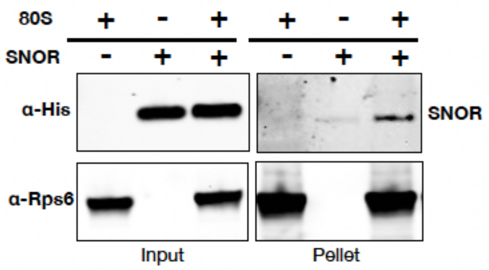


**Extended Data Figure 8. SNOR binds to mammalian ribosomes.**

Immunoblot analysis of different reactions of an in-vitro binding assay between SNOR and ribosomes isolated from HEK293 cells. Input and pellets after centrifugation though a 30% sucrose cushion are shown.

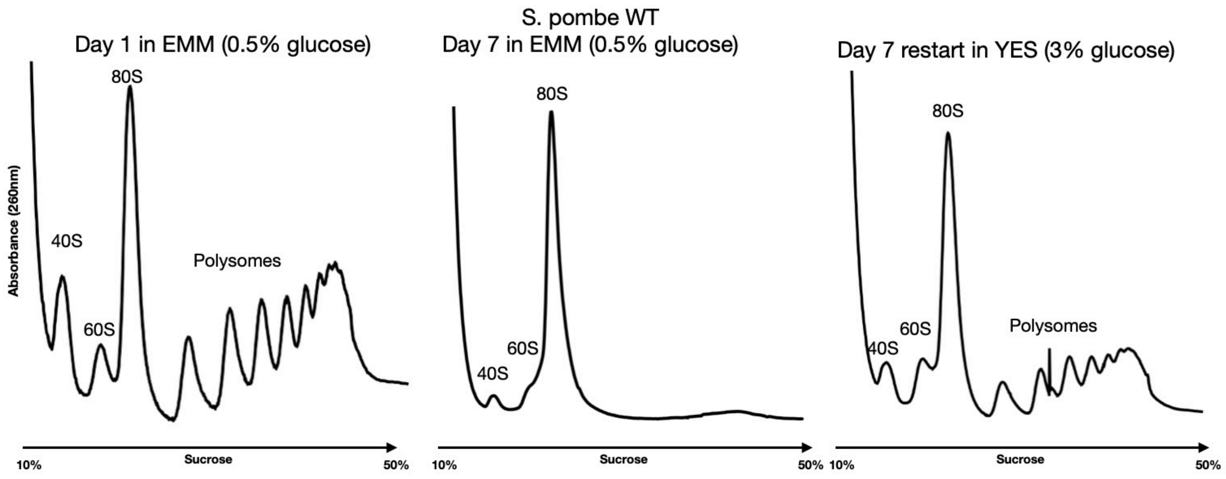


**Extended Data Figure 9. Protein synthesis restart after glucose re-introduction.**

Wildtype *S. pombe* cells (strain 972) were grown in low-glucose EMM (0.5%) for 7 days. Cells were harvested on days 1 and 7, lysed, and subjected to sucrose gradient centrifugation and then fractionated to analyze polysome profiles. By day 7, polysomes shifted to 80S monosomes, indicating a shutdown of protein synthesis, as previously described. Switching the media to YES and incubating the cells for 2 hours led to the reappearance of polysomes, indicating a restart of protein synthesis.

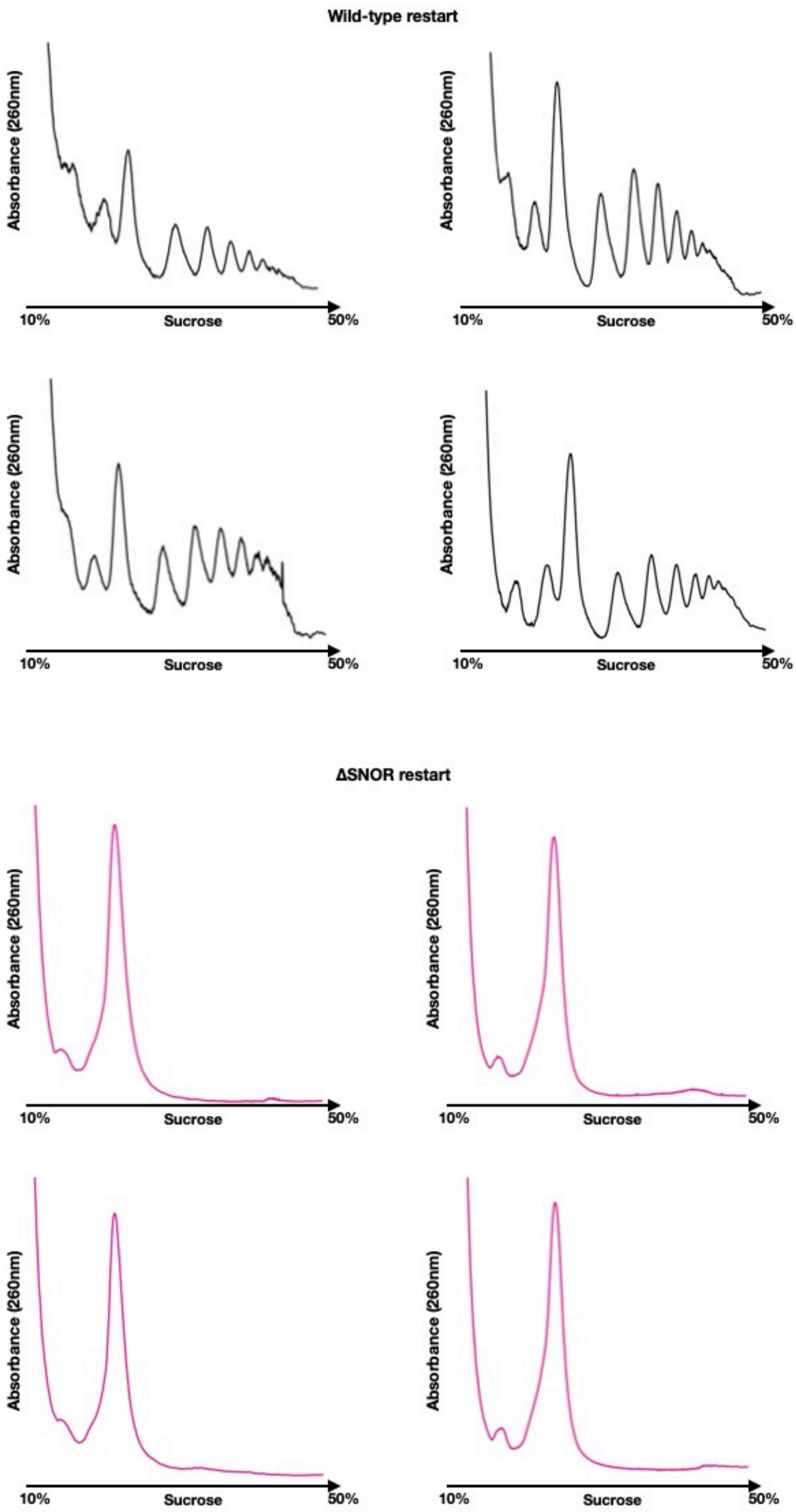
Wild-type restart
Absorbance (260nm)
10%
Sucrose
50%
ΔSNOR restart

**Extended Data Figure 10. SNOR knockout cells fail to restart protein synthesis after glucose depletion.**

Independent biological replicates of polysome gradient profiles from S. pombe cells following glucose reintroduction after 7 days of glucose depletion. Profiles from wildtype cells are shown in black, while profiles from *rtc3*Δ cultures are shown in pink.

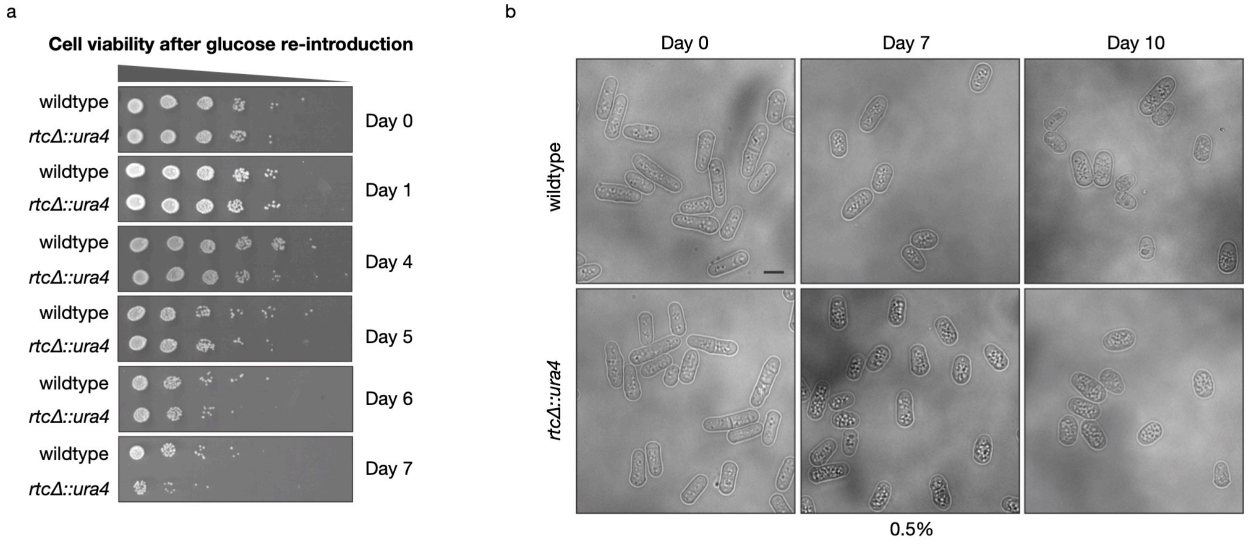


**Extended Data Figure 11.**

(a) Serial dilution spot assays of cultures grown to saturation in EMM with 0.5% glucose at 32˚C and then continuously incubated in the same media shown as a biological duplicate of main Figure 5d. 10-fold serial dilutions of the normalized samples at $OD_{595}$ of 0.25 were spotted on EMM plates containing 0.5% glucose and incubated at 32˚C. Day 1 corresponds to the 24 h time point of continuous incubation. (b) Representative live-cell differential interference contrast images of *wildtype* and *rtc3*Δ cells grown continuously in EMM containing 0.5% glucose for 0, 7 and 10 days.

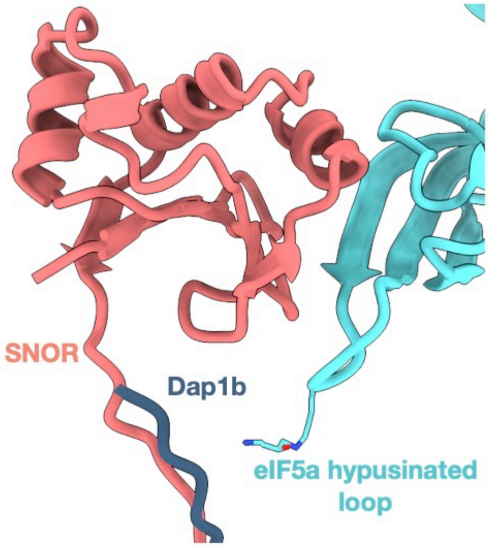


**Extended Data Figure 12.**

**Comparison of the Dap1b binding and the SNOR capping the ribosome exit tunnel.** The hyposinated residues is shown as observed in PDB ID:7OYA.

### Cryo-EM data collection, refinement and validation statistics

| | **Hibernating ribosome (Consensus map)** (EMD- 54290) (PDB 9RVU) | **Hibernating ribosome (OMM-tethered subset)** (EMD-54353) | **Hibernating ribosome (Free cytosolic subset)** (EMD-54354) | **60S:SNOR complex** (EMD-71654) (PDB 9PHC) |
|---|---|---|---|---|
| **Data collection and processing** | | | | |
| Magnification | 64,000x | 64,000x | 64,000x | 105,000x |
| Voltage (kV) | 300 | 300 | 300 | 300 |
| Electron exposure (e–/Å$^2$) | 130 | 130 | 130 | 50 |
| Defocus range (μm) | -2/-6 | -2/-6 | -2/-6 | -1.6/-0.6 |
| Pixel size (Å) | 1.933 | 1.933 | 1.933 | 0.83 |
| Symmetry imposed | | | | |
| Initial particle images (no.) | 43,955 | 20,946 | 23,009 | 502,382 |
| Final particle images (no.) | 26,720 | 12,174 | 14,006 | 53,130 |
| Map resolution (Å) | | | | |
| FSC threshold | 5.5 | 6.0 | 6.7 | 2.91 |
| Map resolution range (Å) | 4.4-12 | | | 2-10 |
| **Refinement** | | | | |
| Initial model used (PDB code) | 9AXV | | | 9AXU |
| Model resolution (Å) | | | | |
| FSC threshold=0.5 (Å) | 5.6 | | | 3.3 |
| Model resolution range (Å) | 4.4-12 | | | 2-10 |
| Map sharpening *B* factor (Å$^2$) | -134.0 | | | 51.2 |
| Model composition | | | | |
| Non-hydrogen atoms | 209,347 | | | 124,378 |
| Protein residues | 12,260 | | | 6,250 |

| | | |
|---|---|---|
| RNA residues | 5,252 | 3,448 |
| *B* factors (Å$^2$) | | |
| Protein | 44.33/346.16/127.39 | 0.00/122.18/45.77 |
| RNA | 46.46/420.12/120.99 | 0.00/184.54/68.22 |
| Ligand | 103.47/197.58/147.50 | 42.19/113.07/88.17 |
| R.m.s. deviations | | |
| Bond lengths (Å) | 0.002 | 0.003 |
| Bond angles (°) | 0.585 | 0.562 |
| Validation | | |
| MolProbity score | 2.17 | 2.04 |
| Clashscore | 16.54 | 7.80 |
| Poor rotamers (%) | 0.00 | 2.68 |
| Ramachandran plot | | |
| Favored (%) | 93.07 | 95.83 |
| Allowed (%) | 6.62 | 4.01 |
| Disallowed (%) | 0.31 | 0.16 |

**Extended Data Table 1. Cryo-ET/Cryo-EM data collection, refinement and validation statistics.**

| Primer | Sequence (5’-3’) |
| --- | --- |
| Act1 Fwd | AAGTACCCCATTGAGCACGG |
| Act1 Rev | TCTCACGGTTGGATTTGGGG |
| Rtc3 Fwd | ATCATCGCCTCAAACGGTCC |
| Rtc3 Rev | TTGTGCTTGCCATGTTCACG |
| Stm1 Fwd | GAAAACCGCTGCTTCTCGTG |
| Stm1 Rev | GCTTCTTTGCCTTCACGAGC |
| eIF5a Fwd | GAACGGCCACGTCGTGATTA |
| eIF5a Rev | GAGCTCACCTTCGGGAAGAC |
| eEF2 Fwd | ACTTGCGTTCTTGCCGTTTC |
| eEF2 Rev | AGAAAACGGGCTCCTGGATG |

**Extended Data Table 2. Real-time qPCR primer sequences.**

| Phylum | Total Genomes | Genomes with SNOR | % Genomes with SNOR |
|---|---|---|---|
| Ascomycota | 1343 | 1264 | 94.1 |
| Basidiomycota | 474 | 412 | 86.9 |
| Blastocladiomycota | 8 | 2 | 25.0 |
| Chytridiomycota | 65 | 35 | 53.8 |
| Cryptomycota | 2 | 1 | 50.0 |
| Microsporidia | 46 | 1 | 2.2 |
| Mucoromycota | 130 | 111 | 85.4 |
| Olpidiomycota | 1 | 0 | 0.0 |
| Zoopagomycota | 179 | 126 | 70.4 |
| Total | 2248 | 1952 | 86.8 |

**Extended Data Table 3. Number of fungal genomes with SNOR across phyla.**

| Query_accession | Sequence |
|---|---|
| SPBC21C3.19.1 | MSSSKANQTRVCYQPEDTTFIIIASNGPDVMRWRKDKTVPLTEIVDSFQVFTTSNNKGNEGQLITASKQELENTFGTSKDVDVVTKILTDGKII |
| NP_011955.1 | MSTVTKYFYKGENTDLIVFAASEELVDEYLKNPSIGKLSEVVELFEVFTPQDGRGAEGELGAASKAQVENEFGKGKKIEEVIDLILRNGKPNSTTSSLKTKGGNAGTKAYN |
| XP_013021238.1 | MSSGPANQTRVVCQTDIASFVVIGASENIIKSWRTDKTIPLTEVVDSFQVFTLTKGSEGELFKASKQQLENAFGTSKDVDVCAKILSEGKISPH |
| XP_013019791.1 | MSSGPANQTRVYYQTDVASFVIIAASEKDVNNWRSDKTIPLTEVVDSFQIFSLNKGSEGELAKASKLELENAFGTSKDVDVCSKILSEGKITPH |
| XP_033766758.1 | MSTVTKYFYKGENTDLIVFATSEELVDEYLKNPSIGKLSEVVEIFEVFTPQDGRGAEGELGAASKAQVENEFGKGKKIEEVIDLILRNGKPNSTTSSLKTKGGNAYK |
| XP_022676094.1 | MSSPIKYFYKGEETDFIIFVNSEEKVQDYLKNSNINNLTEAVSLFKVFANQDARGSEGELGEASKSQIENEFGPKKTTEEVLDLILKNGKPLSS |

**Extended Data Table 4. Query sequences used for HMMER profile building.**

| **Strain** | **Genotype** | **Source** |
|---|---|---|
| AJY19 | YHL 912 h-, ura4-294 leu1-32 | gift Dr. Henry Levin (NIH) |
| AJY23 | *2x-FLAG-rtc3::* ura4-294 leu1-32 | This study |
| KGY28 | *972 h*$^{-}$ | Lab stock |
| KGY6063-2 | *rtc3Δ::ura4*$^{+}$ *h*$^{+}$ | This study |

**Extended Data Table 5 *S. pombe* strains used in this study**
